# Higher-order assembly is a regulatory switch that promotes outer kinetochore recruitment

**DOI:** 10.1101/2023.05.22.541649

**Authors:** Gunter B. Sissoko, Ekaterina V. Tarasovetc, Ekaterina L. Grishchuk, Iain M. Cheeseman

**Affiliations:** Whitehead Institute for Biomedical Research, Cambridge, MA 02142; Department of Biology, Massachusetts Institute of Technology, Cambridge, MA 02142; Department of Physiology, Perelman School of Medicine, University of Pennsylvania, Philadelphia, PA

## Abstract

To faithfully segregate chromosomes during vertebrate mitosis, kinetochore-microtubule interactions must be restricted to a single site on each chromosome. Prior work on pair-wise kinetochore protein interactions has been unable to identify the mechanisms that prevent kinetochore formation in regions with a low density of CENP-A nucleosomes. To investigate the impact of higher-order assembly on kinetochore formation, we generated defined oligomers of the inner kinetochore protein CENP-T using two distinct, genetically engineered systems in human cells. Although individual CENP-T molecules interact poorly with other kinetochore proteins, oligomers that mimic the centromeric density of CENP-T trigger the robust formation of functional, cytoplasmic kinetochore-like particles. Both in cells and *in vitro*, each molecule of oligomerized CENP-T recruits substantially higher levels of outer kinetochore components than monomeric CENP-T molecules. Thus, the density-dependence of CENP-T restricts outer kinetochore recruitment to centromeres, where densely packed CENP-A recruits a high local concentration of CENP-T.

## Introduction

The kinetochore is the essential protein complex that tethers condensed chromosomes to spindle microtubules during mitosis^1, 2^. Individual kinetochore components and subcomplexes have been studied extensively, which has informed our understanding of the mechanisms of kinetochore assembly and function. However, from electron microscopy images, the large copy numbers of proteins at each kinetochore, and recent work on kinetochore protein oligomerization, it has become apparent individual metazoan kinetochores are higher-order assemblies^2–6^. Because previous work has focused primarily on simplified systems, it remains unclear what role higher-order assembly plays in kinetochore biology.

A growing number of proteins are recognized as components of higher-order assemblies with large or undefined stoichiometries like the kinetochore^7–9^. Recent work suggests that the activities of those proteins are modulated by their incorporation into these larger structures, which enables higher-order assembly to spatially regulate cellular activities^7, 9–11^. These structures locally concentrate macromolecules, enabling reactions and interactions within the assemblies that would not otherwise occur at whole-cell concentrations. As the location and number of kinetochores on each chromosome are critical to ensuring proper chromosome segregation and avoiding DNA damage, the spatial regulation conferred by higher-order assembly processes could have important implications for kinetochore formation.

The kinetochore is comprised of two subcomplexes whose assembly mechanisms are tightly controlled in cells. The inner kinetochore is the subset of kinetochore proteins that binds to DNA and localizes to centromeres throughout the cell cycle^1^^2^. Upon mitotic entry, post-translational modifications and the dissolution of the nuclear membrane trigger recruitment of outer kinetochore proteins^12–15^, which perform the kinetochore’s mechanical and signaling functions^12–14,16, 17^. Kinetochore formation is restricted to a single site on each chromosome called the centromere^1, 2^. Additionally, ectopic sites of kinetochore formation result in aberrant chromosome-microtubule interactions, which lead to DNA damage and chromosome segregation errors^18–20^. To direct kinetochore components to centromeres, vertebrate cells mark these regions epigenetically with the histone H3 variant CENP-A^2^^1^. Although CENP-A is necessary to specify the site of kinetochore formation, prior work has found that CENP-A does not drive complete outer kinetochore recruitment when it is incorporated into chromosome arms ^1^^9,2^^2^. Similarly, although complexes of kinetochore proteins have been reconstituted from recombinant proteins *in vitro*^23, 24^, kinetochores do not assemble spontaneously in isolated cytosol^25^. The mechanisms that act alongside CENP-A localization to restrict kinetochore recruitment to centromeres remain unclear. Here, we investigate how human cells confine outer kinetochore recruitment to centromere-localized inner kinetochore assemblies using the emergent properties conferred by higher-order assembly.

To study the role of higher-order assembly in outer kinetochore recruitment, we used artificial oligomers to generate complexes with different valences in cells. We focused on the inner kinetochore protein CENP-T, which directly recruits the outer kinetochore^14, 16, 17, 19^. CENP-T has a structured C-terminal kinetochore localization domain and a disordered N-terminal region with multiple binding sites for outer kinetochore proteins (Figure 1A)^14, 16, 17, 19, 26^. CENP-T is clustered at kinetochores, with approximately 72 copies per human kinetochore^4^. Although CENP-T has no known oligomerization domain, higher-order assembly of the entire inner kinetochore brings CENP-T to a high local concentration^2–4, 6^. Here, we demonstrate that oligomerizing the N-terminal region of CENP-T is sufficient to trigger outer kinetochore recruitment and generate kinetochore-like particles in the cytoplasm. By comparing the interactions of monomeric and oligomeric CENP-T N-termini, we find that CENP-T local concentration regulates its ability to recruit the outer kinetochore, which may restrict complete kinetochore formation to regions with higher-order inner kinetochore assemblies.

**Figure 1:**
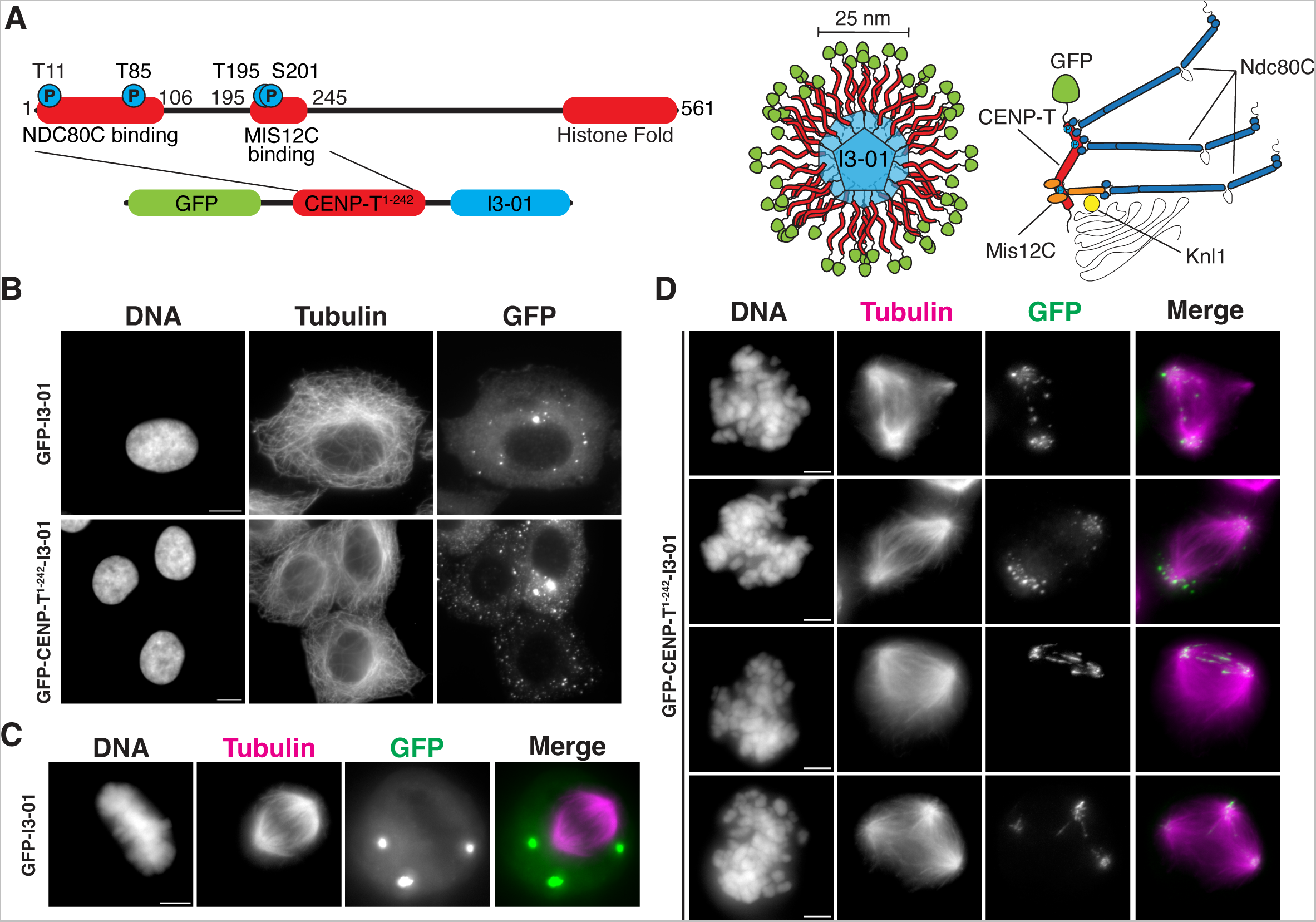
I3-01 oligomerization strategy generates particles that interact with mitotic spindles. (A) Left top: diagram of endogenous CENP-T, its key phosphorylation sites, and the sites of established interactions. Left bottom: construct used to generate CENP-T^1–242^ oligomers in cells. Right: diagrams of the expected oligomers and their predicted interactions with the outer kinetochore. (B) Representative images of CENP-T^1–242^ oligomers and control GFP oligomers in interphase HeLa cells. Scalebars = 10 µm. Different linear image adjustments were used for different fields of view in B-D, so intensities cannot be compared between images. (C) Representative image of control GFP oligomers in mitotic HeLa cells. Scalebars = 5 µm. (D) Examples of CENP-T^1–242^ oligomer localizations in mitotic HeLa cells. Scalebars = 5 µm.

## Results

### Oligomers of the CENP-T N-terminus form kinetochore-like particles

To test the role of higher-order assembly in CENP-T function, we artificially oligomerized the CENP-T N-terminus. A 242 amino acid N-terminal region of CENP-T (CENP-T^1–242^) contains its binding sites for the outer kinetochore complexes NDC80 and MIS12 but lacks its kinetochore localization domain^2, 14, 17, 19, 23, 26^. We fused GFP-CENP-T^1–242^ to I3-01, an oligomerizing tag that forms a 60-subunit homo-oligomer ^27^ (GFP-CENP-T^1–242^–I3-01; Figure 1A). In interphase HeLa cells, GFP-CENP-T^1–242^–I3-01 and GFP-I3-01 control oligomers formed puncta throughout the cytoplasm, consistent with oligomer formation (Figure 1B). In cells with high expression levels, we also observed larger foci, which may reflect a low of level of oligomer aggregation. In mitotic cells, GFP-I3-01 control oligomers also localized throughout the cytoplasm and did not interact with any cellular structures (Figure 1C). By contrast, GFP-CENP-T^1–242^–I3-01 localized consistently to spindle poles and often to spindle microtubules (Figure 1D).

Because CENP-T itself does not bind microtubules, we tested whether GFP-CENP-T^1–242^–I3-01 recruited outer kinetochore components that can interact with the spindle. Using immunofluorescence, we found that the kinetochore’s primary microtubule-binding complex, NDC80, co-localized with GFP-CENP-T^1–242^–I3-01, but not with GFP-I3-01 (Figure 2A). The GFP-CENP-T^1–242^ oligomers also recruited the other core outer kinetochore complexes, MIS12 and KNL1, whereas GFP oligomers did not (Figure 2A; Supplementary Figure 1A). By contrast, the inner kinetochore protein CENP-C did not co-localize with GFP-CENP-T^1–242^–I3-01 (Figure 2A), consistent with prior findings that the CENP-T N-terminus does not interact with other inner kinetochore complexes^19, 26, 28^. To verify these results independently, we immunoprecipitated GFP-CENP-T^1–242^–I3-01 from mitotic HeLa cells. By mass spectrometry, we confirmed that that the NDC80, MIS12, and KNL1 complexes interact with CENP-T^1–242^ oligomers, but not with control oligomers (Figure 2B, Supplementary Figure 1A).

**Figure 2:**
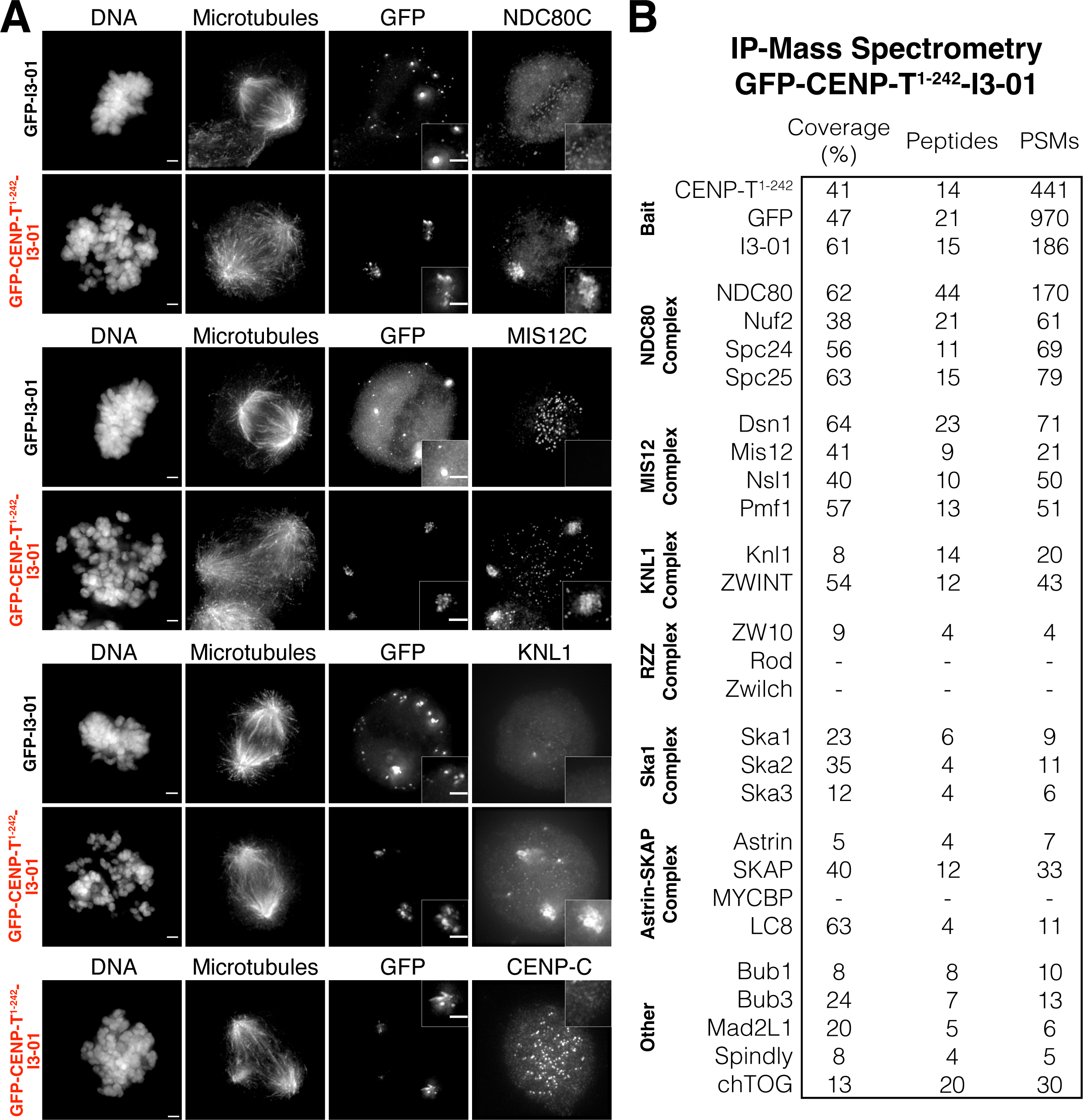
CENP-T^1–242^ oligomers recruit almost the entire outer kinetochore. (A) Co-localization of outer kinetochore proteins with GFP-CENP-T^1–242^-I3-01 oligomers by immunofluorescence. Identical linear image adjustments were used for GFP and kinetochore protein channels for each pair of experimental and control samples. Scalebars = 5 µm. (B) Kinetochore and kinetochore-associated proteins detected in immuno-precipitation mass spectrometry of GFP-CENP-T^1–242^-I3-01.

Downstream of the core outer kinetochore complexes, endogenous kinetochores recruit numerous kinetochore-associated proteins ^29–32^. Among these, the RZZ complex component ZW10 and the SKA1 complex component Ska3 co-localized and immunoprecipitated with GFP-CENP-T^1–242^–I3-01 (Figure 2B, Supplementary Figure 1B). Several other kinetochore-associated proteins, including the Astrin-SKAP complex and components of the spindle assembly checkpoint also co-immunoprecipitated with GFP-CENP-T^1–242^–I3-01 (Figure 2B). These results are remarkable because they suggest that CENP-T^1–242^ recruits a similar set of outer kinetochore components to endogenous kinetochores when oligomerized, generating kinetochore-like particles in the cytoplasm.

### CENP-T-based kinetochore-like particles are functionally similar to endogenous kinetochores

Next, we sought to evaluate the functionality of the GFP-CENP-T^1–242^–I3-01 particles. Kinetochores interact with microtubules in two ways: lateral attachments and end-on attachments^1, 30^. These two modes of interaction enable kinetochores to move processively along microtubules and to track depolymerizing and polymerizing microtubule plus-ends^30, 31, 33–38^. The enrichment of GFP-CENP-T^1–242^–I3-01 at spindle poles and on spindle microtubules suggested that these particles bind to microtubules, but it was unclear whether they engage in the same types of microtubule-driven movement as kinetochores.

To verify that GFP-CENP-T^1–242^–I3-01 particles interact directly with microtubules, we isolated GFP-CENP-T^1–242^–I3-01 from mitotic HeLa cells (Figure 3A; Supplementary figure 2A). Based on the fluorescence intensity of the purified oligomers, they contained 41 ± 5 GFP molecules (Figure 3B). When incubated with stabilized microtubules *in vitro*, the oligomers interacted with the microtubule walls (Figure 3C, D). Using immunofluorescence, we confirmed that the microtubule-bound oligomers had co-purified with the NDC80 complex (Supplementary figure 2C), which likely mediated their interaction with microtubules^2, 39, 40^. By contrast, GFP oligomers isolated from mitotic HeLa cells did not bind microtubules (Figure 3C, D), indicating that microtubule-binding is specific to CENP-T^1–242^ oligomers.

**Figure 3:**
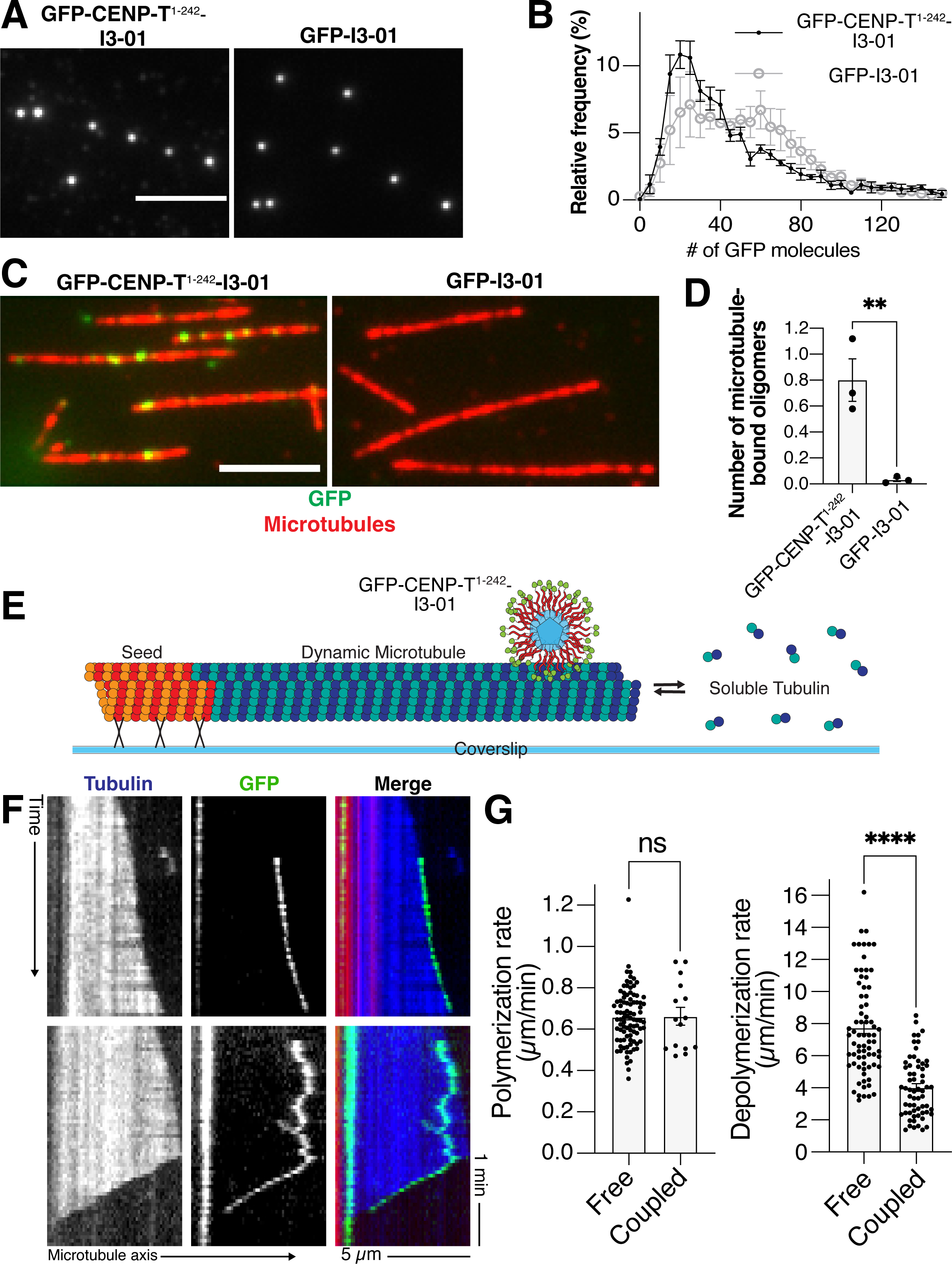
Isolated CENP-T^1–242^ oligomers bind to microtubules and track dynamic microtubule ends. (A) Representative images of GFP-CENP-T^1–242^-I3-01 and GFP-I3-01 isolated from mitotic cells. Scalebar = 5 µm. (B) Histogram showing the distribution of the number of molecules in each oligomer plotted as a percentage of the total number observed of oligomers. Each point represents mean ± SEM from 5 independent experiments, in which more than 180 oligomers were analyzed. Control oligomers contained 51 ± 8 GFP molecules. For more detailed statistics for this and other graphs, see Source data. (C) Representative images of fluorescent microtubules (red) incubated with GFP-tagged CENP-T^1–242^ oligomers and control GFP oligomers (green). Scalebar = 5 µm. (D) Average number of microtubule-bound oligomers in a 10 μm length of microtubule (mean ± SEM from 3 independent experiments). Each point represents an independent experiment in which at least 10 microscopy fields were analyzed. P-values were calculated with unpaired two-tailed t-tests: ** = p<0.01. (E) Schematic of the in vitro assay used to study interactions between CENP-T^1–242^ oligomers and dynamic microtubules. (F) Representative kymographs of dynamic microtubules (tubulin, blue in merge) grown from coverslip-bound microtubule seeds (red in merge) and CENP-T^1–242^ oligomers (GFP, green in merge). Top row: CENP-T^1–242^ oligomer binds directly to polymerizing microtubule end, then processively tracks the end during microtubule polymerization. Bottom row: CENP-T^1–242^ oligomer binds to the wall of a polymerizing microtubule, diffuses on the microtubule lattice, and then tracks the depolymerizing microtubule end. (G) Polymerization and depolymerization rates measured for free microtubule ends and microtubule ends coupled to CENP-T^1–242^ oligomers. Data are based on 3 experiments with particle-free microtubules and 8 experiments with GFP-CENP-T^1–242^-I3-01. Points represent individual microtubule ends; bars show the mean ± SEM. P-values were calculated with unpaired two-tailed t-tests: n.s. = p>0.05; **** = p<0.0001.

To test whether GFP-CENP-T^1–242^ oligomers move on microtubules similarly to kinetochores, we introduced the purified oligomers into chambers containing dynamic microtubules (Figure 3E). Microtubule-bound GFP-CENP-T^1–242^ oligomers exhibited several modes of motility. Some were captured by growing microtubule plus-ends and moved processively with the elongating ends at the rate of tubulin assembly (Figure 3F top, G; Supplementary figure 2D; Supplementary Video 1). Others bound to the microtubule wall, then either remained stationary or moved processively towards the plus-end at 2.7 ± 0.5 µm/min (n=8), a rate that is comparable to that of chromosome congression^41^ (Supplementary figure 2D; Supplementary Video 2). Many microtubule-bound CENP-T^1–242^ oligomers also diffused along the microtubules. Upon encountering a depolymerizing end, these oligomers traveled with the end toward the microtubule seed (Figure 3F bottom, Supplementary figure 2D, Supplementary Video 3). Oligomer-bound ends shortened at half of the rate of oligomer-free ends (Figure 3G). Previous work suggests that mammalian chromosomes and recombinant assemblies of human CENP-T, NDC80, and MIS12 cause a similar suppression of microtubule depolymerization ^42, 43^ Together, these results suggest that CENP-T^1–242^ oligomerization triggers the formation of outer kinetochore structures that interact with microtubules similarly to endogenous kinetochores. Thus, the localization of GFP-CENP-T^1–242^–I3-01 oligomers to spindle poles may reflect depolymerization-driven movement toward microtubule minus-ends in the absence of polar ejection forces and attachments to sister kinetochores, which normally antagonize poleward movement.

### Oligomerized CENP-T recruits outer kinetochore proteins more efficiently than CENP-T monomers

The ability of GFP-CENP-T^1–242^ oligomers to form kinetochore-like particles in the cytoplasm is surprising because our prior work found virtually no interactions between soluble CENP-T and the NDC80 complex in mitotic HeLa cell extract^25^. To investigate how the GFP-CENP-T^1–242^ oligomers differ from monomers, we directly compared the behaviors of GFP-CENP-T^1–242^ oligomers with those of identical GFP-CENP-T^1–242^ constructs without an oligomerizing tag (GFP-CENP-T^1–242^).

Unlike CENP-T^1–242^ oligomers, which localized to mitotic spindles in immunofluorescence experiments, GFP-CENP-T^1–242^ monomers dispersed diffusely throughout the cytoplasms of mitotic cells and interacted minimally with spindles (Figure 1C, 4A). The more robust spindle localization of oligomers could reflect increased microtubule-binding avidity relative to GFP-CENP-T^1–242^ monomers^33, 35, 43, 44^. It could also result from improved outer kinetochore recruitment when GFP-CENP-T^1–242^ is oligomerized. If the latter is true, we predicted that GFP-CENP-T^1–242^ oligomers would compete more effectively with endogenous kinetochores for outer kinetochore components than monomers expressed at comparable levels, resulting in distinct phenotypes.

**Figure 4:**
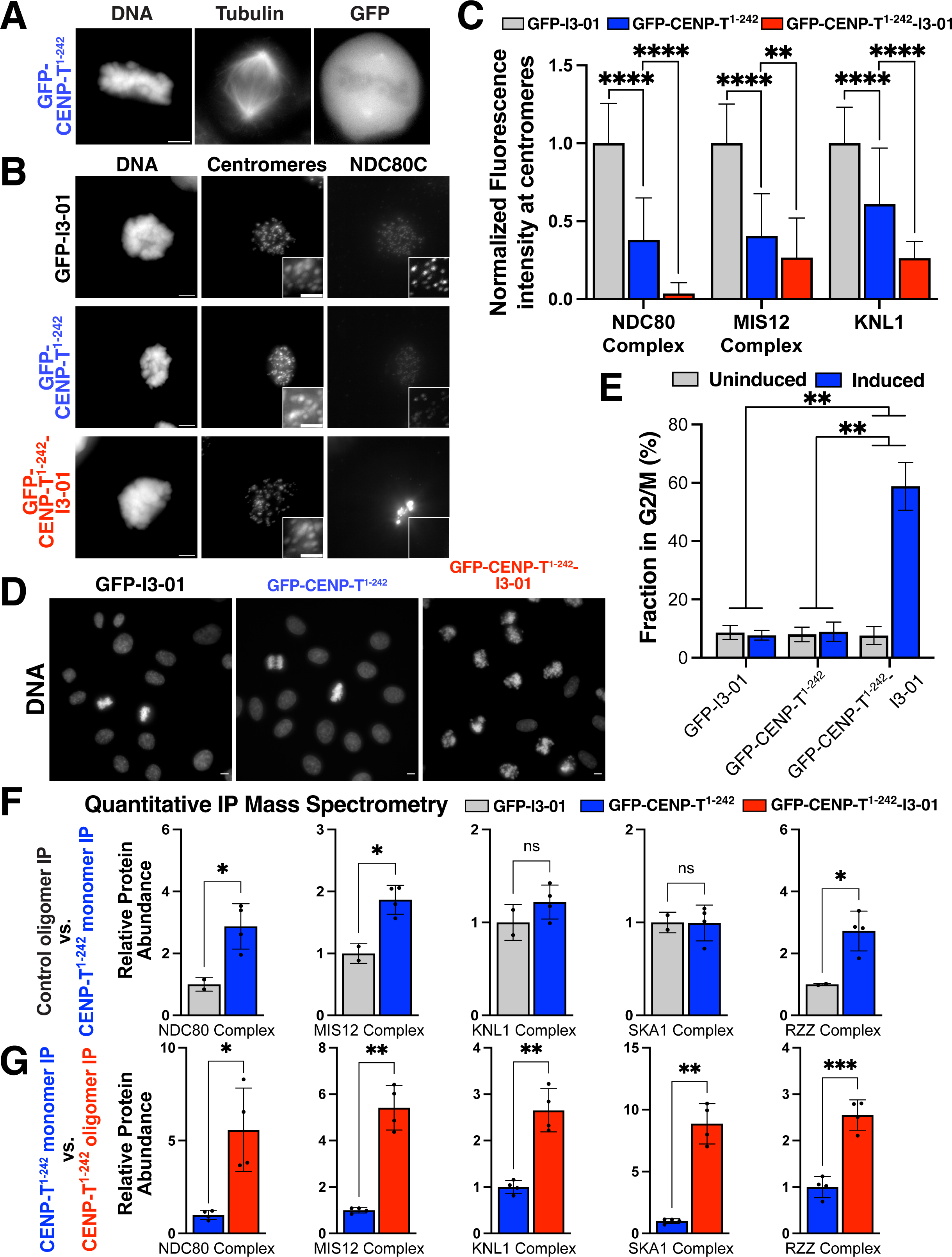
CENP-T^1–242^ oligomers recruit the outer kinetochore more robustly than monomeric CENP-T^1–242^. (A) Representative image of monomeric CENP-T^1–242^ in mitotic HeLa cells. (B) Representative immunofluorescence images of NDC80 levels at centromeres in cell expressing control oligomers, monomeric CENP-T^1–242^, and CENP-T^1–242^ oligomers. All cells are mitotically arrested with S-Trityl-L-Cysteine. Identical linear image adjustments were used for all conditions in centromere and NDC80 complex channels. NDC80 insets are adjusted differently from full-size images. Full-sized image scalebars = 5 µm. Inset scalebars = 2 µm. (C) Quantification of NDC80 complex levels from (A) and MIS12 complex and KNL1 levels in similar experiments. Each bar represents the mean ± SD of outer kinetochore protein signal from cells expressing the designated construct. Values for each cell were calculated from the mean of the outer kinetochore protein signals of kinetochore in that cell. Before calculating the mean for a cell, the outer kinetochore protein signal of each kinetochore in the cell was normalized to anti-centromere antibody signal from that kinetochore. Overall means were calculated from pooled data from multiple experiments. To make results comparable between experiments, the mean for each cell was normalized to the mean of all cells in the GFP-I3-01 control sample in the same experiment. NDC80 complex: 30 cells for each condition from 2 experiments. Mis12 complex: 42-61 cells for each condition from 3 experiments. KNL1: 27-34 cells for each condition from 2 experiments. P-values were calculated using Welch’s two-tailed t-test. ** = p <0.01; **** = p <0.0001. (D) Representative images of fields of cells stained with Hoechst expressing control GFP-I3-01 oligomers, GFP-CENP-T^1–242^ monomers, or GFP-CENP-T^1–242^ oligomers. Scalebar = 5 µm. (E) Percentage of cells in G2/M as determined by DNA content measurements by flow cytometry in cell lines expressing control GFP oligomers, GFP-CENP-T^1–242^ monomers, or GFP-CENP-T^1–242^ oligomers. Expression of the constructs was controlled by a TetOn system and induced with Doxycycline. Bars represent mean percentage ± SD from 3 repeats. Statistical analysis was performed on the differences between means of induced and uninduced. P-values were calculated using Welch’s two-tailed t-test. ** = p<0.01. (F) Comparison of outer kinetochore protein co-immuno-precipitation by control oligomers and monomeric CENP-T^1–242^ by TMT-based quantitative immuno-precipitation mass spectrometry. Each point represents a biological replicate from a single multiplexed mass spectrometry run. Raw abundances of proteins in each sample were normalized to the abundance of GFP peptides, then the normalized abundances for the designated complexes were expressed as multiples of the normalized abundance in the GFP-I3-01 sample. Complex abundances were obtained by calculating the sum of abundances of the complex components. P-values were calculated using Welch’s two-tailed t-test. * = p < 0.05. (G) Comparison of outer kinetochore protein co-immuno-precipitation by CENP-T^1–242^ oligomers and CENP-T^1–242^ monomers by TMT-based quantitative immune-precipitation (IP) mass spectrometry. Each point represents a biological replicate from a single multiplexed mass spectrometry run. Raw abundances were processed as described in (F), but normalized to the abundance of GFP-CENP-T^1–242^, and normalized abundances for the designated proteins or complexes were expressed as multiples of the normalized abundance in the GFP-CENP-T^1–242^ sample. Complex abundances were obtained by calculating the sum of abundances of the complex components. P-values were calculated using Welch’s two-tailed t-test. * = p < 0.05; ** = p <0.01; *** = p < 0.001.

To test how GFP-CENP-T^1–242^ expression impacted endogenous kinetochores, we measured the localization of core outer kinetochore complexes to centromeres in cells arrested in mitosis and expressing GFP-CENP-T^1–242^ monomers, GFP-CENP-T^1–242^ oligomers, or GFP oligomers. Expression of monomeric GFP-CENP-T^1–242^ had a moderate effect on outer kinetochore recruitment, with NDC80 levels at centromeres reduced to 38% of the levels control cells expressing GFP oligomers. By contrast, expression of comparable levels of GFP-CENP-T^1–242^–I3-01 severely depleted outer kinetochore proteins from endogenous kinetochores (Figure 4B, C; Supplementary figure 3A, B). This was particularly true for the NDC80 complex, which was reduced to 3.7% of control levels in cells expressing GFP-CENP-T^1–242^–I3-01 (Figure 4B, C).

The recruitment of the outer kinetochore to exogenous oligomers and the resulting depletion of core kinetochore proteins from endogenous kinetochores had a dramatic effect on mitotic progression. Expression of GFP-CENP-T^1–242^–I3-01, but not monomeric CENP-T^1–242^, led to severe mitotic defects, including misaligned chromosomes, spindle abnormalities, and a potent mitotic arrest (Figures 1D, 2A, 4D). After 24 hours of GFP-CENP-T^1–242^–I3-01 expression, the fraction of cells in G2/M increased from 6.7% to 64.5% by DNA content analysis (Figure 4E), whereas monomeric GFP-CENP-T^1–242^ expression had no impact on the fraction of cells in G2/M (Figure 4E). Because both forms of GFP-CENP-T^1–242^ were expressed at similar levels (Supplementary Figure 3A, B), the systemic impacts of GFP-CENP-T^1–242^ oligomerization on the cells suggest that oligomerization enables CENP-T N-termini to recruit the outer kinetochore components more efficiently.

To measure each construct’s outer kinetochore recruitment efficiency directly, we immunoprecipitated CENP-T^1–242^ monomers, CENP-T^1–242^ oligomers, and control oligomers from HeLa cells arrested in mitosis, and compared the abundances of interacting partners using tandem mass tag-based quantitative mass spectrometry. To enable a direct comparison, we normalized all protein abundances to the abundance of peptides shared between pairs of bait proteins. For example, in comparisons between GFP-CENP-T^1–242^–I3-01 and GFP-CENP-T^1–242^, outer kinetochore protein abundances were normalized to the abundance of GFP-CENP-T^1–242^ peptides. Using this approach, we determined that monomeric GFP-CENP-T^1–242^ co-purified with more NDC80 complex and MIS12 complex than the GFP-I3-01 control (Figure 4F), consistent with a modest interaction between monomeric CENP-T^1–242^ and outer kinetochore components. Strikingly, GFP-CENP-T^1–242^–I3-01 oligomers associated with substantially larger amounts of core outer kinetochore proteins than monomeric GFP-CENP-T^1–242^, with 5.6-fold more NDC80 complex per CENP-T molecule, 5.4-fold more MIS12 complex, and 2.7-fold more KNL1 complex (Figure 4G). Furthermore, GFP-CENP-T^1–242^–I3-01 recruited higher levels of downstream outer kinetochore and kinetochore-associated proteins, including the SKA1 complex, the RZZ complex, Spindly, Mad2L1, and chTOG (Figure 4G, Supplementary figure 4B). Thus, when CENP-T^1–242^ is oligomerized, each molecule recruits outer kinetochore proteins more efficiently, explaining why CENP-T oligomers form functional kinetochore-like particles in cells and impair mitotic progression.

### Increasing the size of CENP-T oligomers incrementally improves recruitment of outer kinetochore components

The ability of artificial CENP-T oligomers to compete with endogenous kinetochores and produce kinetochore-like particles in mitotic cells strongly suggests that high local concentrations of CENP-T activate outer kinetochore recruitment. However, the precise oligomer size required to achieve this effect was unclear. To determine whether the oligomerization-dependent recruitment has an oligomer size threshold or gradually activates as GFP-CENP-T^1–242^ oligomer size increases, we used an unrelated strategy called the “SunTag” to precisely manipulate the stoichiometry of CENP-T^1–242^ oligomers ^45^. The SunTag is a two-component system with a single-chain monoclonal antibody (scFv), which we fused to CENP-T^1–242^ (scFv-sfGFP-CENP-T^1–242^), and a scaffold with multiple repeats of the antibody’s cognate epitope (GCN4pep; Figure 5A). When scFv-sfGFP-CENP-T^1–242^ is co-expressed with the scaffold, one copy of the scFv-sfGFP-CENP-T^1–242^ fusion protein can bind to each GCN4pep repeat, resulting in oligomers of defined sizes^45^ (Figure 5A). We ensured equal scFv-sfGFP-CENP-T^1–242^ expression levels by generating all cell lines from the same scFv-sfGFP-CENP-T^1–242^–expressing parental line (Supplementary figure 5A, B, E, G).

**Figure 5:**
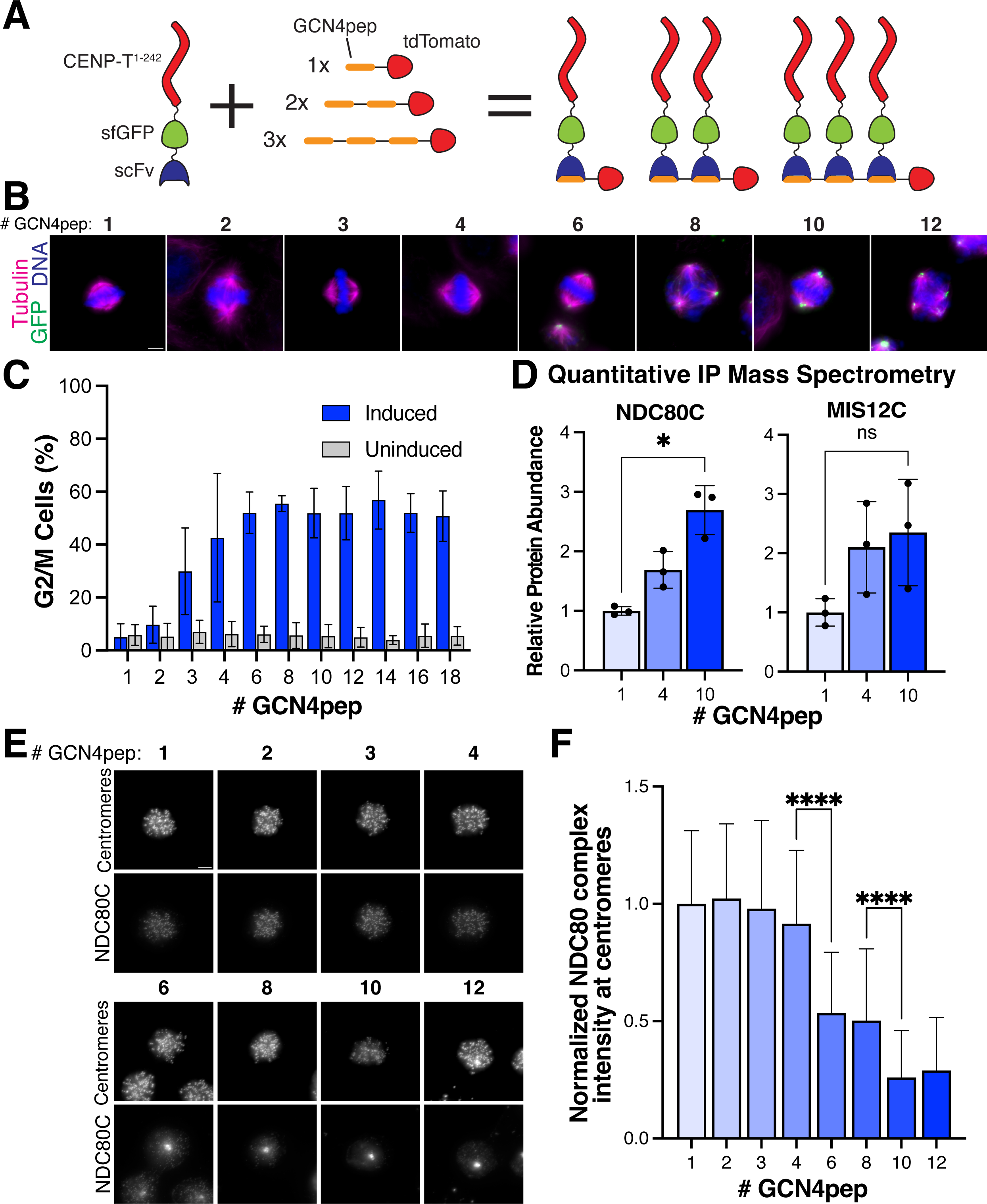
Each additional CENP-T^1–242^ molecule incrementally increases outer kinetochore recruitment of neighboring molecules. (A) Diagram of SunTag tunable oligomerization strategy. (B) Representative immunofluorescence images of SunTag oligomer localization with different numbers of GCN4pep on the scaffold. GFP signal in all images is scaled the same. Scalebar = 5 µm. (C) Percentage of cells in G2/M as determined from DNA content measurements by flow cytometry in cell lines expressing SunTag with scaffolds with different numbers of GCN4pep. scFv-sfGFP-CENP-T^1–242^ expression was controlled by a TetOn system and induced with doxycycline. Bars represent mean percentage of cells in G2/M ± SD from 3 repeats. Welch’s ANOVA test was performed on the differences between means of induced and uninduced to calculate a P-value, p < 0.0001. (D) Comparison of outer kinetochore protein co-immuno-precipitation by scFv-sfGFP-CENP-T^1–242^ expressed with scaffolds with 1, 4, or 10 copies of GCN4pep by TMT-based quantitative immuno-precipitation mass spectrometry. TdTomato-tagged scaffolds were immuno-precipitated, then abundances for isolated proteins were normalized to the abundance of scFv-sfGFP-CENP-T^1–242^ peptides. Normalized abundances for the designated complexes were expressed as multiples of the abundance in the 1xGCN4pep sample. Complex abundances were obtained by calculating the sum of abundances of the complex components. Each point represents a biological replicate from a single multiplexed mass spectrometry run. P-values were calculated using Welch’s two-tailed t-test. * = p < 0.05; n.s. = p > 0.05. (E) Representative immunofluorescence images of NDC80 levels at centromeres in cell expressing the scFv-sfGFP-CENP-T^1–242^ with scaffolds with different numbers of GCN4pep. All cells are mitotically arrested with S-Trityl-L-Cysteine. All images use the same linear image adjustments. Scalebar = 5 µm. (F) Quantification of NDC80 complex levels from (E). Each bar represents the mean ± SD of NDC80 signal from cells expressing the designated construct. Overall means were calculated from pooled data from multiple experiments. To make results comparable between experiments, the mean for each cell was normalized to the mean of all cells in the 1xGCN4pep sample in the same experiment. 40-46 total cells were analyzed from 3 different experiments for each condition. Welch’s ANOVA test was performed to calculate P-value for the whole dataset (p < 0.0001). Welch’s t test was used to calculated P-value for selected pairwise comparisons. **** = p < 0.0001.

When co-expressed with a single GCN4pep repeat (1xGCN4pep), scFv-sfGFP-CENP-T^1–242^ did not localize to the mitotic spindle, like monomeric GFP-CENP-T^1–242^ (Figures 1B, 5B). As we increased the number of GCN4pep repeats, we observed mitotic abnormalities and sfGFP-scFv-CENP-T^1–242^ began to localize to spindle poles. With 6 or more GCN4pep repeats, sfGFP-scFv-CENP-T^1–242^ robustly localized to spindle poles and spindle microtubules, like GFP-CENP-T^1–242^–I3-01 kinetochore particles, which have on average 40 molecules of CENP-T^1–242^ (Figure 3B, 5B). Similarly, scFv-sfGFP-CENP-T^1–242^ expression with 1xGCN4pep or 2xGCN4pep did not cause a cell cycle arrest, but the fraction of cells in G2/M increased gradually from 2 to 6 GCN4 repeats. Larger oligomers caused a potent mitotic arrest with 40-60% of cells in G2/M (Figure 5C). Thus, CENP-T^1–242^’s ability to interact with spindle microtubules and impair mitotic progression increases incrementally as additional molecules are added to an oligomer, and the maximum phenotypic effect can be elicited with 6-8 CENP-T N-termini.

We expected that larger CENP-T oligomers generated with the SunTag system would be similar to I3-01 oligomers: they would recruit outer kinetochore components and compete with the endogenous kinetochores. However, it was unclear whether intermediately sized oligomers would behave similarly to large oligomers or exhibit an intermediate behavior. To answer this question, we performed quantitative IP-mass spectrometry on scFv-sfGFP-CENP-T^1–242^ co-expressed with scaffolds with either 1, 4, or 10 GCN4pep repeats (1xGCN4pep, 4xGCN4pep, 10xGCN4pep). We immunoprecipitated the tdTomato-tagged scaffolds to isolate only the scaffold-bound scFv-sfGFP-CENP-T^1–242^ molecules. We normalized the abundances of co-immunoprecipitated proteins in each sample to the abundance of scFv-sfGFP-CENP-T^1–242^ in the sample to determine the relative amount of protein recruited per molecule of scFv-sfGFP-CENP-T^1–242^. The NDC80 complex was 1.7-fold enriched in the immunoprecipitation of 4xGCN4pep relative to the immunoprecipitation of 1xGCN4pep (Figure 5D). In the 10xGCN4pep immunoprecipitation, NDC80 was further enriched to 2.7-fold its abundance in the 1xGCN4pep immunoprecipitation (Figure 5D). In addition to the NDC80 complex, we observed a gradual increase in the co-immunoprecipitation of other outer kinetochore components as we increased the number of GCN4pep repeats, although these results were not statistically significant (Figure 5D; Supplementary figure 5H). Consistent with this gradual increase in outer kinetochore recruitment, as the number of binding sites on the scaffold increased, we observed a corresponding reduction in the levels of the NDC80 and MIS12 complexes at centromeres (Figure 5E, F; Supplementary figure 5I), which indicates that larger oligomers stripped more outer kinetochore proteins from endogenous kinetochores than smaller oligomers. Together, these results suggest that outer kinetochore recruitment by a GFP-CENP-T^1–242^ molecule in an oligomer increases incrementally as additional molecules are added to the oligomer. In addition, they show that an orthologous oligomerization system can recapitulate the effect of the I3-01 oligomerization system on CENP-T, which confirms that the enhancement of outer kinetochore recruitment is the result of oligomerization, not an effect specific to I3-01.

### Oligomerized CENP-T uses known binding sites to recruit NDC80

CENP-T has two known binding sites for the NDC80 complex that are activated by phosphorylation at T11 and T85 and required for outer kinetochore recruitment at endogenous kinetochores^13, 14, 16, 17, 19^. To confirm that outer kinetochore recruitment by oligomeric CENP-T^1–242^ uses the same pathways, we tested whether the enhancement of NDC80 recruitment upon oligomerization is dependent on T11 and T85 phosphorylation. In the SunTag system, we mutated T11 and T85 to alanine to prevent phosphorylation (scFv-sfGFP-CENP-T^1–242/2TA^). When we expressed scFv-sfGFP-CENP-T^1–242/2TA^ in cells, NDC80 levels at endogenous kinetochores were comparable for all scaffold sizes (Supplementary Figure 5K). This suggests that CENP-T^1–242^ oligomers use the previously characterized binding sites to recruit NDC80, consistent with our conclusion that the assembled complexes are *bona fide* kinetochore particles.

### Oligomerization of CENP-T is necessary to saturate NDC80-binding sites

Our experiments in mitotic HeLa cells revealed that oligomerization of CENP-T enables robust recruitment of outer kinetochore proteins. In cells, the underlying mechanisms could be complex and involve additional factors such as post-translational modifications or microtubule interactions. To determine if additional components are involved in enhancing CENP-T outer kinetochore recruitment, we reconstituted the interaction between the NDC80 complex and the CENP-T N-terminus *in vitro*. To activate NDC80 recruitment, we used CENP-T^1–242^ constructs with the phosphomimetic substitutions T11D, T27D, and T85D (GFP-CENP-T^1–242/3D^) (Figure 6A). We expressed and purified GFP-I3-01 control oligomers, GFP-CENP-T^1–242/3D^-I3-01 oligomers, and GFP-CENP-T^1–, 242/3D^ monomers from *E. coli*, and we visualized them using total internal reflection fluorescence microscopy. When imaged under identical conditions, purified GFP-CENP-T^1–242/3D^-I3-01 complexes appeared much brighter than the GFP-CENP-T^1–242/3D^ monomers (Figure 6B), as expected. By normalizing the intensities of these fluorescent foci to the intensity of a single GFP fluorophore, we determined that recombinant GFP-CENP-T^1–242/3D^-I3-01 complexes contain 66 ± 10 GFP-CENP-T^1–242/3D^ molecules, consistent with oligomer formation (Supplementary figure 7A).

**Figure 6:**
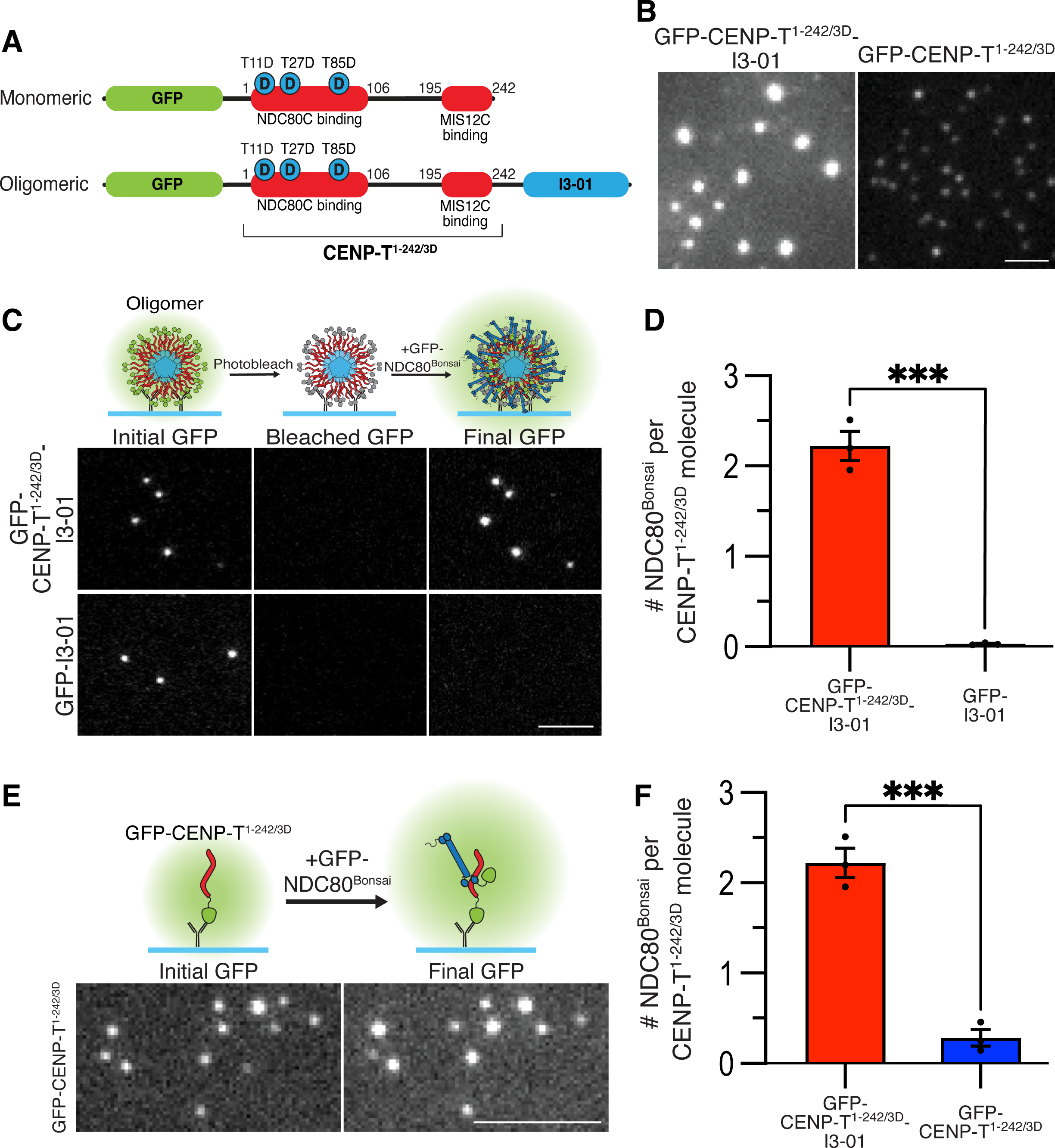
Oligomerization of CENP-T activates NDC80 recruitment in vitro. (A) Diagram of GFP-CENP-T^1–242/3D^-I3-01 (top) and GFP-CENP-T^1–242/3D^ (bottom) constructs. Both constructs contain CENP-T^1–242^ region with activating phospho-mimetic substitutions at sites T11, T27, and T85. (B) Representative images of GFP-CENP-T^1–242/3D^-I3-01 oligomers or GFP-CENP-T^1–242/3D^ monomers taken with identical microscope settings and shown with identical linear image adjustments. Scalebar = 2 µm. (C) Top: Diagram of single molecule experimental approach with GFP-CENP-T^1–242/3D^-I3-01 oligomers immobilized on coverslips. The initial GFP intensity of each oligomer, which represented the number of CENP-T^1–242/3D^ molecules per oligomer, was recorded (initial GFP). Before addition of GFP-tagged NDC80^Bonsai^, oligomers were photobleached to make recruitment of GFP-tagged NDC80^Bonsai^ more apparent. After addition of GFP-NDC80^Bonsai^, the GFP signal was measured again (Final GFP). Final GFP represents the number of NDC80^Bonsai^ molecules bound, which allowed us to determine CENP-T-NDC80^Bonsai^ binding efficiency. Bottom: representative images of GFP-CENP-T^1–242/3TD^-I3-01 and GFP-I3-01 oligomers immobilized on coverslips before photobleaching, after photobleaching, and after interaction with 100 nM GFP-tagged NDC80^Bonsai^. Scalebar = 5 µm. (D) Efficiency of NDC80 recruitment to GFP-CENP-T^1–242/3D^-I3-01, and control GFP-I3-01. The result is the number of NDC80 molecules bound per molecule in an oligomer. Bars are mean ± SEM from 3 independent experiments. Each point is the median result from independent trials with at least 12 oligomers analyzed. P-values were calculated by unpaired two-tailed t-test: *** = p<0.001. (E) Top: diagram of single molecule experimental approach with GFP-CENP-T^1–242/3D^ monomers. The GFP signal from each monomer was recorded before addition of GFP-NDC80^Bonsai^ (Initial GFP). After addition of GFP-NDC80^Bonsai^, the GFP signal was measured again (Final GFP). Bottom: representative examples of GFP-CENP-T^1–242/3D^ monomers before and after interaction with 100 nM GFP-tagged NDC80^Bonsai^. Scalebar = 5 µm. (F) Efficiency of NDC80 recruitment to GFP-CENP-T^1–242/3D^-I3-01 oligomers and GFP-CENP-T^1–242/3TD^ monomers. Bars are mean ± SEM from 3 independent experiments. Each point is the median result from independent trials with at least 12 oligomers or 33 monomers analyzed. Data for GFP-CENP-T^1–242/3D^-I3-01 oligomer is duplicated from (D). Each point is the average result from an independent trial, bars are mean ± SEM. P-values were calculated by unpaired two-tailed t-test: *** = p<0.001.

**Figure 7:**
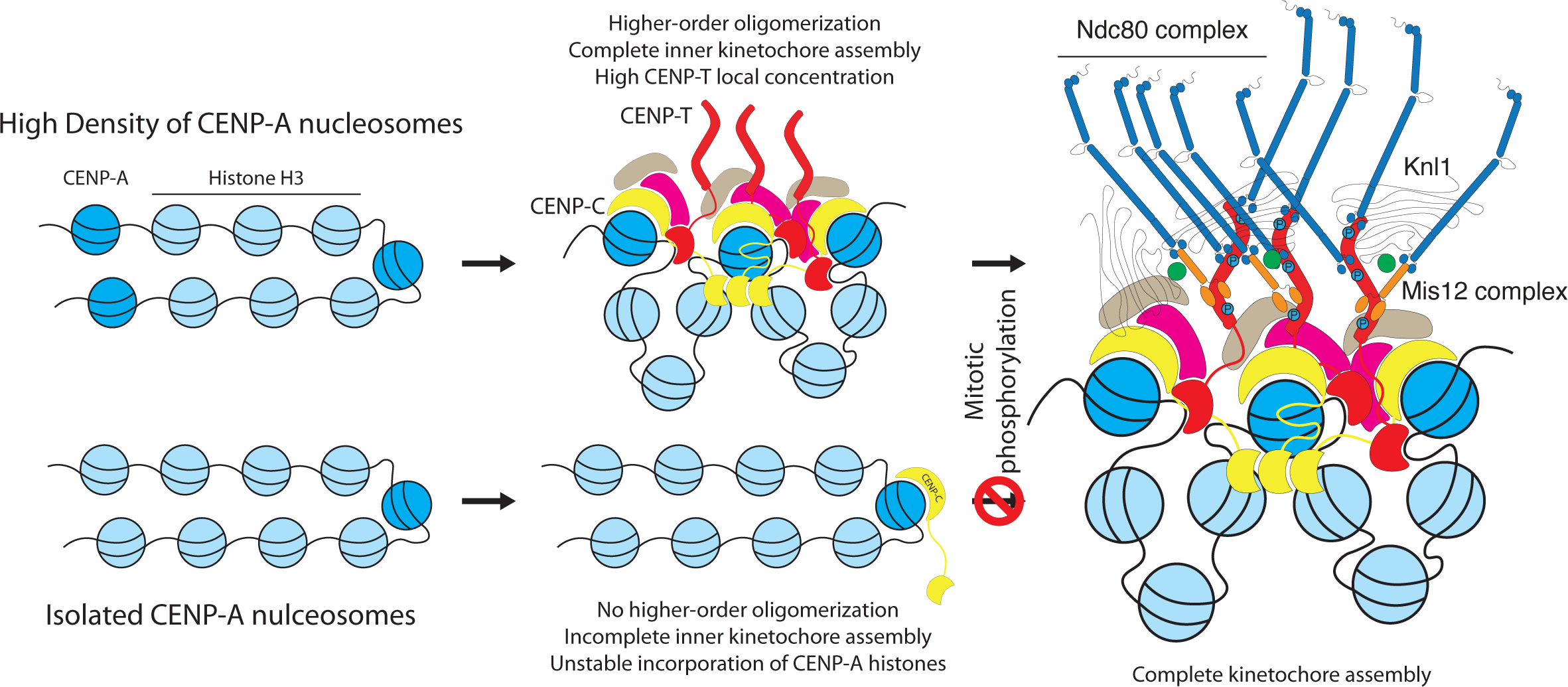
Model of the role of higher order oligomerization in kinetochore assembly. In regions where CENP-A nucleosomes are at a high density (>25% of nucleosomes; top of figure), they recruit CENP-C, which can use its Cupin domain to form CENP-A-CENP-C oligomers. Those oligomers can then recruit multiple inner kinetochore modules, which protect CENP-A from eviction when replication forks pass through during S-phase. Those stable inner kinetochore oligomers have a high local concentration of CENP-T, which enables CENP-T to robustly recruit the outer kinetochore when it is phosphorylated by Cdk1 during mitosis, generating complete kinetochores. When CENP-A is deposited at a low density (bottom), it may be able to recruit individual molecules of CENP-C, but it cannot generate inner kinetochore oligomers. As a result, is evicted during S phase and any CENP-A that remains fails to recruit the rest of the kinetochore.

To determine how efficiently NDC80 binds to oligomers, we immobilized oligomers on a coverslip and measured the GFP signal from individual foci. This initial GFP intensity corresponded to the number of GFP molecules per focus. We then photobleached the oligomers and incubated them with recombinant GFP-tagged NDC80^Bonsai^ complex, a shortened version of the NDC80 complex ^46^ (Figure 6C). For these experiments, we used 100 nM NDC80^Bonsai^, which is close to the reported concentration of NDC80 in human cells^47, 48^. After a 10-minute incubation, we washed away unbound NDC80^Bonsai^ and measured the GFP signal from each focus again. This final GFP intensity corresponded to the number of NDC80^Bonsai^ molecules per focus (Figure 6C). By normalizing the final GFP signal of each focus to its initial signal, we determined the number of NDC80 complexes bound to each GFP or GFP-CENP-T^1–, 242/3D^ molecule in an oligomer. With this approach, we found that each oligomerized molecule of GFP-CENP-T^1–242/3D^ bound to 2.2 ± 0.2 NDC80^Bonsai^ molecules (Figure 6D, Supplementary figure 7F). As the CENP-T N-terminus has two direct NDC80-binding sites, this result indicates that each CENP-T molecule in an oligomer saturates its NDC80-binding sites ^4, 14, 17, 23^. GFP control oligomers did not bind to NDC80^Bonsai^ (Figure 6D, Supplementary figure 7F), as expected. Furthermore, NDC80^ΔSpc^^24/25^ (also known as NDC80^Broccoli^), which lacks the CENP-T-binding region of the complex^16, 49^, failed to interact with GFP-CENP-T^1–242/3D^ oligomers (Supplementary figure 7D-F), confirming that the observed interaction depends on the known CENP-T-NDC80 binding interfaces. To compare the CENP-T^1–242/3D^ oligomers to monomers, we performed analogous experiments on monomeric CENP-T^1–242/3D^ without the photobleaching step (Figure 6E). Strikingly, each GFP-CENP-T^1–242/3TD^ molecule only recruited 0.28 ± 0.09 NDC80^Bonsai^ complexes (Figure 6F). Because binding events are binary, this result means that most monomeric GFP-CENP-T^1–242/3D^ molecules did not bind any NDC80^Bonsai^. To ensure that this result was not due to photobleaching of the monomer between the initial and final GFP measurements, we measured the fraction of molecules photobleached as a function of exposure time. In the 0.6 second exposure time used in this experiment, a negligible 4% of GFP molecules were photobleached (Supplementary figure 7G). Thus, CENP-T^1–242/3D^ must be oligomerized to saturate its direct NDC80-binding sites at physiological NDC80 concentrations. As our single molecule system lacks any other factors found in cells, these experiments also demonstrate that oligomerization-dependence is an intrinsic feature of the CENP-T-NDC80 interaction and does not depend on interactions with other factors, such as microtubules. Additionally, because both CENP-T^1–242^ oligomers and monomers had phosphomimetic mutations, these results show that CENP-T oligomerization is a regulatory mechanism downstream of activation by mitotic phosphorylation and that the change in NDC80 binding is not mediated by changes in CENP-T phosphorylation.

## Discussion

### Higher-order oligomerization dictates the site of functional kinetochore formation

Previous work has established the hierarchy of kinetochore recruitment to centromeres but has not explored how the higher-order organization of kinetochores contributes to their assembly. Based on prior work *in vitro* and *in silico*, the large copy numbers of proteins at individual kinetochores were thought to be necessary to form dynamic load-bearing microtubule attachments^43, 50^. Here, we show that this density is also an important regulatory cue that may restrict kinetochore formation to centromeres. By mimicking endogenous kinetochore organization through artificial oligomerization of the inner kinetochore protein CENP-T *in vivo* and *in vitro*, we found that CENP-T only interacts stably with outer kinetochore proteins when it is oligomerized. We observed this behavior with two unrelated oligomerization methods, which shows that the local concentration of CENP-T molecules is the important regulatory factor. Our results demonstrate that the CENP-T-NDC80 interaction is intrinsically dependent on CENP-T local concentration and indicate that each molecule of CENP-T recruits a gradually increasing amount of outer kinetochore components as neighboring CENP-T molecules are added.

These findings suggest that higher-order assembly has an important role in centromere specification and spatially restricting the site of kinetochore formation. Centromeres are specified epigenetically by histone H3-variant CENP-A^18, 51^. However, the incorporation of CENP-A into chromatin is not sufficient to trigger complete kinetochore formation^19, 22, 52, 53^. Our work supports a model in which higher-order assembly regulates the location of kinetochores by restricting outer kinetochore recruitment to regions with a high density of inner kinetochore complexes. In this model, dense deposition of CENP-A at centromeres is coupled to active oligomerization of the inner kinetochore by proteins such as CENP-C or CENP-N^3, 6, 54, 55^. Clustering of repetitive inner kinetochore structures generates a high local concentration of CENP-T, which primes the inner kinetochore for outer kinetochore recruitment during mitosis (Figure 7). In addition, it is possible that higher-order oligomerization of the inner kinetochore has additional roles in initiating kinetochore assembly, such as triggering inner kinetochore recruitment to densely deposited CENP-A nucleosomes and protecting CENP-A nucleosomes from eviction during S phase ^52^ (Figure 7). This paradigm for controlling kinetochore formation is complementary to previously defined regulation, such as post-translational modifications. Our *in vitro* experiments with phospho-mimetic CENP-T mutants suggest that, even when kinase activity is permissive, the dependence on higher-order assembly prevents aberrant formation of functional outer kinetochore complexes at non-centromeric sites on chromosomes and on cytoplasmic kinetochore components that have not been incorporated into the chromatin^25^.

### Artificial kinetochore-like particles as *in vitro* tools for biophysical analysis of human kinetochores

*In vitro* biophysical analysis of kinetochores depends on tools that can recapitulate endogenous kinetochore-microtubule interactions outside of cells. In budding yeast, which have much smaller point centromeres and simplified kinetochores, it has been possible to isolate intact kinetochore complexes for *in vitro* analysis^56–59^. However, the more complex vertebrate kinetochore is far less tractable, so biophysical work on human kinetochores has been limited to simplified systems such as the NDC80 complex alone or assemblies of the NDC80 and SKA1 complexes^33, 35, 43, 44, 60, 61^. We previously generated ectopic kinetochores by targeting the CENP-T N-terminus to chromosomal *lacO* arrays. In that system, CENP-T-based ectopic kinetochores were functional and rescued excision of endogenous kinetochores^19, 62^. In this work, we used a related strategy to generate kinetochore-like particles that can be purified. We demonstrate that these particles are compositionally and mechanically similar to endogenous human kinetochores, and that they are sufficiently tractable for *in vitro* applications. These properties suggest that CENP-T-based particles can be used for *in vitro* biophysical assays to yield insights into the mechanical properties of complete human kinetochores.

### Identifying the mechanism by which local concentration regulates CENP-T activity

Recent interest in higher-order protein assemblies has focused on liquid-liquid phase separation as a mechanism for locally concentrating interacting partners^10^. These membrane-less compartments are thought to form through the interactions of proteins with disordered regions that contain multivalent low-affinity interfaces with dissociation constants in the micromolar or millimolar ranges^9, 56, 57^. Such biomolecular condensates may use platforms such as membranes or nucleic acids to nucleate their formation by bringing together many copies of disordered scaffolds^9, 58^. Like those proteins, CENP-T is multivalent and disordered^2, 14, 17^. However, unlike putative phase-separating scaffolds, CENP-T uses high affinity binding sites with dissociation constants in the nanomolar range to interact with outer kinetochore proteins^15, 16, 23^. Furthermore, the transition from a homogenous mixture to a phase separated solution is binary and happens when a phase-separating scaffold achieves its saturation concentration^10^, which is not consistent with the gradual change in binding that we observed with the SunTag system. As a result, existing models of phase separation are unlikely to explain our findings. This work suggests that CENP-T makes use of high local concentrations in a manner that is dependent on specific high-affinity interactions that enable stable binding.

### Applying artificial oligomerization approaches to other pathways

In addition to kinetochores, other biological pathways are thought to use higher-order assemblies to regulate their activities. For example, numerous signal transduction pathways form massive complexes called “signalosomes” to initiate intracellular signaling^9, 59–62^. Establishing direct links between oligomerization and function has been one of the challenges of studying these higher-order assemblies. The toolkit that we have used could prove valuable for investigating these relationships. Unlike popular higher-order oligomerization systems such as the optogenetic CRY2 system, the I3-01 and SunTag oligomerization systems generate stable and tunable oligomers, respectively ^27, 45, 63, 65^. These oligomers can be purified for *in vitro* applications and used to study the stoichiometries that govern the activities of higher-order assemblies. Our approach is readily applicable to other proteins that have distinct oligomerization and functional domains, both at the kinetochore and in unrelated pathways.

## Materials and Methods

### Plasmid cloning

The I3-01 gene was synthesized by Genewiz. sfGFP-scFv tag and CENP-T^1–1242/2TA^ were synthesized by Twist Bioscience. SunTag scaffolds were obtained from pcDNA4TO-mito-mCherry-24xGCN4_v1, which was a gift from Ron Vale (Addgene plasmid #60913). CENP-T^1–242^ was obtained from pKG174^19^. Lentiviral plasmids were generated from Lenti-Cas9-2A-Blast, which was a gift from Jason Moffat (Addgene plasmid #73310).

### Cell line generation

The cell lines used in this study are described in Table 1. Doxycycline-inducible cell lines were generated by homology-directed insertion into the AAVS1 “safe-harbor” locus. Donor plasmid containing selection marker, the tetracycline-responsive promoter, the transgene, and reverse tetracycline-controlled transactivator flanked by AAVS1 homology arms^64^ was transfected using Lipofectamine 2000 with a pX330-based plasmid^66^ expressing both spCas9 and a guide RNA specific for the AAVS1 locus (pNM220; gRNA sequence – 5’-GGGGCCACTAGGGACAGGAT). Cells were selected with 0.5 µg/mL puromycin (Life Technologies). Clonal lines were obtained by fluorescence activated cell-sorting single cells into 96 well plates.

**Table 1:** Cell lines used in this study.

| **Cell Lines** | **Description** | **Clonality** | **Expression** | **Source** |
| --- | --- | --- | --- | --- |
| **cGS49** | EGFP-I3-01 | Polyclonal | Inducible | pGS96 integrated at AAVS safe-harbor locus |
| **cGS50** | EGFP-CENP-T<sup>1-242</sup>-I3-01 | Clonal | Inducible | pGS97 integrated at AAVS safe-harbor locus |
| **cGS54** | EGFP- CENP-T<sup>1-242</sup> | Clonal | Inducible | pGS101 integrated at AAVS safe-harbor locus |
| **cGS107** | TdTomatoLAP-10xGCN4 w/ scFv-sfGFP-CENP-T<sup>1-242</sup> | Clonal | Scaffold constitutive, scFv inducible | pGS241 integrated at AAVS safe-harbor locus; lentiviral transduction of pGS174 |
| **cGS115** | TdTomatoLAP-1xGCN4 w/ scFv-sfGFP-CENP-T<sup>1-242</sup> | Clonal | Scaffold constitutive, scFv inducible | pGS241 integrated at AAVS safe-harbor locus; lentiviral transduction of pGS281 |
| **cGS117** | TdTomatoLAP-3xGCN4 w/ scFv-sfGFP-CENP-T<sup>1-242</sup> | Clonal | Scaffold constitutive, scFv inducible | pGS241 integrated at AAVS safe-harbor locus; lentiviral transduction of pGS283 |
| **cGS365** | tdTomato-2xGCN4-T2A-BSD w/ scFv-sfGFP- CENP-T<sup>1-242</sup> | Polyclonal | Scaffold constitutive, scFv inducible | pGS241 integrated at AAVS safe-harbor locus; lentiviral transduction of pGS374 |
| **cGS366** | tdTomato-3xGCN4-T2A-BSD w/ scFv-sfGFP- CENP-T<sup>1-242</sup> | Polyclonal | Scaffold constitutive, scFv inducible | pGS241 integrated at AAVS safe-harbor locus; lentiviral transduction of pGS375 |
| **cGS367** | tdTomato-4xGCN4-T2A-BSD w/scFv-sfGFP- CENP-T<sup>1-242</sup> | Polyclonal | Scaffold constitutive, scFv inducible | pGS241 integrated at AAVS safe-harbor locus; lentiviral transduction of pGS376 |
| **cGS368** | tdTomato-6xGCN4-T2A-BSD w/ scFv-sfGFP- CENP-T<sup>1-242</sup> | Polyclonal | Scaffold constitutive, scFv inducible | pGS241 integrated at AAVS safe-harbor locus; lentiviral transduction of pGS377 |
| **cGS369** | tdTomato-8xGCN4-T2A-BSD w/ scFv-sfGFP- CENP-T<sup>1-242</sup> | Polyclonal | Scaffold constitutive, scFv inducible | pGS241 integrated at AAVS safe-harbor locus; lentiviral transduction of pGS378 |
| **cGS370** | tdTomato-10xGCN4-T2A-BSD w/ scFv-sfGFP- CENP-T<sup>1-242</sup> | Polyclonal | Scaffold constitutive, scFv inducible | pGS241 integrated at AAVS safe-harbor locus; lentiviral transduction of pGS379 |
| **cGS371** | tdTomato-12xGCN4-T2A-BSD w/ scFv-sfGFP- CENP-T<sup>1-242</sup> | Polyclonal | Scaffold constitutive, scFv inducible | pGS241 integrated at AAVS safe-harbor locus; lentiviral transduction of pGS380 |
| **cGS372** | tdTomato-14xGCN4-T2A-BSD w/ scFv-sfGFP- CENP-T<sup>1-242</sup> | Polyclonal | Scaffold constitutive, scFv inducible | pGS241 integrated at AAVS safe-harbor locus; lentiviral transduction of pGS381 |
| **cGS373** | tdTomato-16xGCN4-T2A-BSD w/ scFv-sfGFP- CENP-T<sup>1-242</sup> | Polyclonal | Scaffold constitutive, scFv inducible | pGS241 integrated at AAVS safe-harbor locus; lentiviral transduction of pGS382 |
| **cGS374** | tdTomato-18xGCN4-T2A-BSD w/ scFv-sfGFP- CENP-T<sup>1-242</sup> | Polyclonal | Scaffold constitutive, scFv inducible | pGS241 integrated at AAVS safe-harbor locus; lentiviral transduction of pGS383 |
| **cGS386** | tdTomato-1xGCN4-T2A-BSD w/ scFv-sfGFP- CENP-T<sup>1-242</sup> | Polyclonal | Scaffold constitutive, scFv inducible | pGS241 integrated at AAVS safe-harbor locus; lentiviral transduction of pGS373 |
| **cGS416** | myc-1xGCN4-T2A-BSD w/ scFv-sfGFP- CENP-T<sup>1-242</sup> | Polyclonal | Scaffold constitutive, scFv inducible | pGS241 integrated at AAVS safe-harbor locus; lentiviral transduction of pGS395 |
| **cGS417** | myc-2xGCN4-T2A-BSD w/ scFv-sfGFP- CENP-T<sup>1-242</sup> | Polyclonal | Scaffold constitutive, scFv inducible | pGS241 integrated at AAVS safe-harbor locus; lentiviral transduction of pGS396 |
| **cGS418** | myc-3xGCN4-T2A-BSD w/<br>scFv-sfGFP-CENP-T<sup>1-242</sup> | Polyclonal | Scaffold constitutive,<br>scFv inducible | pGS241 integrated at AAVS safe-harbor locus; lentiviral transduction of pGS397 |
| **cGS419** | myc-4xGCN4-T2A-BSD w/<br>scFv-sfGFP-CENP-T<sup>1-242</sup> | Polyclonal | Scaffold constitutive,<br>scFv inducible | pGS241 integrated at AAVS safe-harbor locus; lentiviral transduction of pGS398 |
| **cGS420** | myc-6xGCN4-T2A-BSD w/<br>scFv-sfGFP-CENP-T<sup>1-242</sup> | Polyclonal | Scaffold constitutive,<br>scFv inducible | pGS241 integrated at AAVS safe-harbor locus; lentiviral transduction of pGS399 |
| **cGS421** | myc-8xGCN4-T2A-BSD w/<br>scFv-sfGFP-CENP-T<sup>1-242</sup> | Polyclonal | Scaffold constitutive,<br>scFv inducible | pGS241 integrated at AAVS safe-harbor locus; lentiviral transduction of pGS400 |
| **cGS422** | myc-10xGCN4-T2A-BSD w/<br>scFv-sfGFP-CENP-T<sup>1-242</sup> | Polyclonal | Scaffold constitutive,<br>scFv inducible | pGS241 integrated at AAVS safe-harbor locus; lentiviral transduction of pGS401 |
| **cGS423** | myc-12xGCN4-T2A-BSD w/<br>scFv-sfGFP-CENP-T<sup>1-242</sup> | Polyclonal | Scaffold constitutive,<br>scFv inducible | pGS241 integrated at AAVS safe-harbor locus; lentiviral transduction of pGS402 |
| **cGS642** | myc-1xGCN4-T2A-BSD w/<br>scFv-sfGFP-CENP-T<sup>1-242/2TA</sup> | Polyclonal | Scaffold constitutive,<br>scFv inducible | pGS403 integrated at AAVS safe-harbor locus; lentiviral transduction of pGS395 |
| **cGS643** | myc-2xGCN4-T2A-BSD w/<br>scFv-sfGFP-CENP-T<sup>1-242/2TA</sup> | Polyclonal | Scaffold constitutive,<br>scFv inducible | pGS403 integrated at AAVS safe-harbor locus; lentiviral transduction of pGS396 |
| **cGS644** | myc-3xGCN4-T2A-BSD w/<br>scFv-sfGFP-CENP-T<sup>1-242/2TA</sup> | Polyclonal | Scaffold constitutive,<br>scFv inducible | pGS403 integrated at AAVS safe-harbor locus; lentiviral transduction of pGS397 |
| **cGS645** | myc-4xGCN4-T2A-BSD w/<br>scFv-sfGFP-CENP-T<sup>1-242/2TA</sup> | Polyclonal | Scaffold constitutive,<br>scFv inducible | pGS403 integrated at AAVS safe-harbor locus; lentiviral transduction of pGS398 |
| **cGS646** | myc-6xGCN4-T2A-BSD w/<br>scFv-sfGFP-CENP-T<sup>1-242/2TA</sup> | Polyclonal | Scaffold constitutive,<br>scFv inducible | pGS403 integrated at AAVS safe-harbor locus; lentiviral transduction of pGS399 |
| **cGS647** | myc-8xGCN4-T2A-BSD w/scFv-sfGFP-CENP-T<sup>1-242/2TA</sup> | Polyclonal | Scaffold constitutive, scFv inducible | pGS403 integrated at AAVS safe-harbor locus; lentiviral transduction of pGS400 |
| **cGS648** | myc-10xGCN4-T2A-BSD w/scFv-sfGFP-CENP-T<sup>1-242/2TA</sup> | Polyclonal | Scaffold constitutive, scFv inducible | pGS403 integrated at AAVS safe-harbor locus; lentiviral transduction of pGS401 |
| **cGS649** | myc-12xGCN4-T2A-BSD w/scFv-sfGFP-CENP-T<sup>1-242/2TA</sup> | Polyclonal | Scaffold constitutive, scFv inducible | pGS403 integrated at AAVS safe-harbor locus; lentiviral transduction of pGS402 |

Cell lines containing SunTag scaffolds were generated by lentiviral transduction. Lentivirus was generated by using Xtremegene-9 (Roche 06365787001) to co-transfect the scaffold-containing pLenti plasmid, VSV-G envelope plasmid, and Delta-VPR or psPAX2 (gift from Didier Trono; Addgene plasmid #12260) packaging plasmids into HEK-293T cells^67^. cGS107, cGS115, and cGS117 were sorted for tdTomato-positive cells. Other lentivirus cell lines were selected with 2 µg/mL blasticidin (Life Technologies). Cell lines containing SunTag scaffolds were generated from clonal parental lines expressing the desired sfGFP-scFv construct at comparable levels.

### Cell Culture

HeLa cells were cultured in Dulbecco’s modified Eagle medium (DMEM) supplemented with 10% fetal bovine serum, 100 U/mL penicillin and streptomycin, and 2 mM L-glutamine at 37°C with 5% CO_2_. TetOn cell lines were cultured in FBS certified as tetracycline-free. TetOn constructs were induced with 1 μg/mL doxycycline for 24 hours. To depolymerize microtubules, cells were treated with 3.3 μM Nocodazole for 16 hours. To arrest cells in mitosis, cells were treated with 10 μM STLC for 16 hours. Hela cells were regularly monitored for mycoplasma contamination.

### Western blot

Cells were harvested by trypsinization and resuspended, then washed with PBS and immediately lysed on ice for 30 min in fresh urea lysis buffer (50 mM Tris pH 7.5, 150 mM NaCl, 0.5% NP-40, 0.1% SDS, 6.5 M Urea, 1X Complete EDTA-free protease inhibitor cocktail (Roche), 1 mM PMSF) or cells lysed directly on plate with RIPA buffer (150 mM NaCl, 1% Nonident P-40 substitute, 0.5% Sodium Deoxycholate, 0.1% Sodium Dodecyl Sulfate, 50 mM Tris pH 7.5, 1X Complete EDTA-free protease inhibitor cocktail (Roche), 1 mM PMSF) on ice. Protein concentrations were measured using either Bradford reagent (Bio-Rad) or BCA protein assay kit (Pierce) and used to normalize loading. Antibodies are listed in Table 2. Primary antibodies were diluted in Blocking Buffer and applied to the membrane for 1 hour. HRP-conjugated secondary antibodies (GE Healthcare; Digital) were diluted in TBST (TBS with 0.1% Tween-20). Clarity enhanced chemiluminescence substrate (Bio-Rad) was used according to the manufacturer’s instructions. Membranes were imaged with a KwikQuant Imager (Kindle Biosciences). HRP was quenched by agitation in 0.2% Sodium Azide in TBST for at least 1 hour.

**Table 2:** Primary Antibodies used in this study.

| Antibody | Source | Use | Identifier |
| --- | --- | --- | --- |
| α-Tubulin | Sigma-Aldrich | 1:2500 (IF) | <u>CAT#: T9026-.5ML;</u><br><u>RRID: AB_477593</u> |
| Tubulin Beta 3 | Serotec |  | MCA2047 |
| S-tag | Abcam |  | Ab87840 |
| MIS12 Complex (Human) | Cheeseman lab<sup>75</sup> | 1:1000 (IF) | Available from MilliporeSigma, CAT# ABE2585 |
| KNL1 (Human) | Cheeseman lab<sup>76</sup> | 1:1000 (IF) | N/A |
| Ndc80-Bonsai (Human) | Cheeseman lab<sup>49</sup> | 1:4800 (WB, IF) | N/A |
| CENP-C (Human) | Cheeseman lab<sup>19</sup> | 1:16000 (IF) | N/A |
| ZW10 (Human) | Abcam | 1:1000 (IF) | ab21582 |
| Ska3/Rama1 (Human) | Cheeseman lab<sup>77</sup> | 1:1000 (IF) | N/A |
| Anti-centromere Antibodies (Human) | Antibodies, Inc. | 1:100 (IF) | CAT#: 15-234-0001 |
| GFP | Cheeseman lab | IP | N/A |
| mCherry | Cheeseman lab | IP | N/A |
| GFP | Roche | 1:5000 (WB) | CAT#11814460001 |
| β-actin (Human) (HRP conjugated) | Fisher Scientific | 1:5000 (WB) | #MA515739HRP |
| T2A | Sigma-Aldrich | 1:1000 (WB) | #MABE1923-100UG |

### Immunofluorescence and microscopy of mitotic cells

Cells were seeded on poly-L-lysine (Sigma-Aldrich) coated coverslips and fixed as indicated in Table 3. Coverslips were washed with 0.1% PBS-Tx (PBS with 0.1% Triton X-100) and blocked in Abdil (20 mM Tris-HCl, 150 mM NaCl, 0.1% Triton X-100, 3% bovine serum albumin, 0.1% NaN_3_, pH 7.5). Primary antibodies used in this study are described in Table 2 and were diluted in Abdil. Dilutions are listed in Table 2 (IF = immunofluorescence, WB = Western blot, IP = immunoprecipitation). Cy3- and Cy5-conjugated (or Alexa 647-conjugated) secondary antibodies (Jackson ImmunoResearch Laboratories) were diluted 1:300 in 0.1% PBS-Tx. DNA was stained with 1 µg/mL Hoechst-33342 (Sigma-Aldrich) in 0.1% PBS-Tx for 10 min. Coverslips were mounted with PPDM (0.5% *p*-phenylenediamine, 20 mM Tris-HCl, pH 8.8, 90% glycerol). Images were acquired with a DeltaVision Ultra High-Resolution microscope (Imsol) and deconvolved where indicated. All images are maximal intensity projections in z. Image analysis was performed in Fiji (ImageJ, NIH)^68^.

**Table 3:** Immunofluorescence fixation conditions.

| Figure | Fixation Conditions |
| --- | --- |
| 1B-D, 2A, 4A, S1A, 4D | 4% formaldehyde in PHEM 10min |
| 4B, 5E | 0.25% PBS-Tx pre-extraction 5min followed by 4% formaldehyde in 0.125% PBS-Tx 10 min |
| 5B | 0.25% PHEM-Tx pre-extraction 5min followed by 4% formaldehyde in 0.125% PHEM-Tx 10 min |

Integrated fluorescence intensity of mitotic centromeres was measured with a custom CellProfiler 4.0 pipeline^69^ (adapted from McQuin, et al. 2014^70^). The median intensity of a 5-pixel region surrounding each centromere was multiplied by the area of the centromere to determine background intensity. Regions with high GFP signal were masked to avoid measuring kinetochore proteins bound to GFP-tagged constructs. Values for each cell were calculated from the mean of the outer kinetochore protein antibody signals of kinetochores in that cell. Before calculating the mean for a cell, the kinetochore protein antibody intensity of each kinetochore in the cell was normalized to anti-centromere antibody signal from that kinetochore.

### DNA content analysis

Cells were incubated in 1μg/mL doxycycline for 24 hours. 5 mM EDTA, 20 μg/mL Hoechst-33342 (Sigma-Aldrich), and 10 μM Verapimil (Tocris; Spirochrome) were added directly to media for 30 minutes to 1 hour to detached cells from the plate and stain them. Cells were collected and filtered through 35 μm nylon mesh (Falcon). Hoechst, GFP, and tdTomato signals were measured on an LSRFortessa (BD Biosciences) flow cytometer. Results were analyzed with FlowJo software. Example gating strategy for SunTag system is shown in Supplementary figure 6. The fraction of cells in each cell cycle phase was determined in FlowJo with a Watson (Pragmatic) model using the Cell Cycle tool. The DNA content of at least 9,000 cells was analyzed for each condition per experiment.

### Crosslinking Immunoprecipitation-Mass Spectrometry

Construct expression was induced, and cells were arrest in mitosis as described in the cell culture section. They were harvested 24 hours after doxycycline addition and 16 hours after STLC addition by mitotic shake off. Mitotic cells were centrifuged at 250 RCF and resuspended in Crosslinking Buffer (20 mM Hepes pH 7.5, 10 mM KCl, 1.5 mM MgCl_2_, 1 mM DTT). To crosslink samples, formaldehyde was added to 0.1% and samples were incubated at 37°C for 15 minutes. Glycine was added to 0.25 M to quench formaldehyde. Samples were washed once in PBS and once in Lysis Buffer without detergent (25 mM Hepes pH 8.0, 2 mM MgCl_2_, 0.1 mM EDTA pH 8.0, 0.5 mM EGTA pH 8.0, 150 mM KCl, 15% Glycerol).

To prepare protein extracts, samples were thawed and an equal volume of 1.5X high salt lysis buffer w/ detergent (37.5 mM Hepes pH 8.0, 3 mM MgCl_2_, 0.15 mM EDTA pH 8.0, 0.75 mM EGTA pH 8.0, 450 mM KCl, 15% Glycerol, 0.225% Nonidet P-40 substitute) was added. Proteases were inhibited with a tablet of Complete EDTA-free protease inhibitor cocktail (Roche) and 1mM PMSF. Phosphatases were inhibited with 0.4 mM Sodium Orthovanadate, 5 mM Sodium Fluoride, and 20 mM Beta-glycerophosphate. Cells were lysed with Branson Digital Sonifier tip sonicator to shear DNA. Lysates were for incubated 1 hour at room temperature with Protein A beads (Bio-Rad) coupled to anti-GFP antibodies (Cheeseman lab). After incubation, beads were washed in Lysis buffer with high salt, DTT, and LPC (25 mM Hepes pH 8.0, 2 mM MgCl_2_, 0.1 mM EDTA pH 8.0, 0.5 mM EGTA pH 8.0, 300 mM KCl, 15% Glycerol, 0.15% Nonidet P-40 subsitute, 1 mM DTT, 10 μg/mL leupeptin (Millipore), 10 μg/mL pepsin (Thermo Fisher Scientific), 10 μg/mL chymostatin (Millipore)), then washed in the same buffer without detergent once (25 mM Hepes pH 8.0, 2 mM MgCl_2_, 0.1 mM EDTA pH 8.0, 0.5 mM EGTA pH 8.0, 300 mM KCl, 15% Glycerol, 1 mM DTT, 10 μg/mL leupeptin (Millipore), 10 μg/mL pepsin (Thermo Fisher Scientific), 10 μg/mL chymostatin (Millipore)). Beads were incubated in 0.1 M glycine pH 2.6 for 5 minutes 3 times and once with Lysis buffer without detergent to elute. Elutions were pooled and Tris pH 8.5 was added to 200 mM. Eluate was incubated at 65°C for 1.5 hours to reverse crosslinks. Proteins were precipitated with 20% Trichloroacetic Acid (Fisher Bioreagents) overnight on ice. The next day, samples were centrifuged at 20817 RCF at 4°C. Pellets were washed twice with ice cold Acetone. Samples were dried in Eppendorf Vacufuge and stored at -80°C.

Samples were resuspended in SDS lysis buffer (5%, 50 mM TEAB pH 8.5), then DTT was added to 20 mM and samples were incubated at 95°C for 10 minutes. After cooling to room temperature, samples were treated 40 mM iodoacetamide (Sigma) for 30 minutes in the dark. Samples were acidified with 1.2% phosphoric acid, then run over S-Trap microcolumns (ProtiFi), digested on the columns, and eluted as described in ProtiFi S-trap micro kit protocol. We quantified eluate peptide concentration with Quantitative Fluorometric Peptide Assay (Pierce). We lyophilized remaining eluate to remove solvent and stored at -80°C.

For quantitative mass spectrometry, up to 10 samples were prepared simultaneously as described above. Each sample was incubated with a different TMT10plex label (Thermo Fisher Scientific) in 30% acetonitrile, 24.5 mM TEAB pH 8.5 for 1 hour at room temperature. TMT10plex reagents were added to labeling reactions in a 10-fold excess over peptides by mass. The labeling reaction was quenched by adding hydroxylamine to 0.3% and incubating for 15 minutes at room temperature. Labeled samples were pooled, then lyophilized to remove solvent, and stored at -80°C.

To removed salt and labels and to increase coverage, samples were fractionated with High pH Reversed-Phase Peptide Fractionation kit (Pierce). After fractionation, fractions were lyophilized and resuspended in 0.1% formic acid. Samples were analyzed on an Orbitrap Exploris 480 connected to an EASY-nLC chromatography system using two compensation voltages applied with a FAIMS Pro Interface (Thermo Fisher Scientific). Proteins were identified in Proteome Discoverer 2.4 (Thermo Fisher Scientific) using Sequest HT. Peptide-spectrum matches were validated using Percolator. Tandem Mass Tag quantification was done in Proteome Discoverer. For quantitative mass spectrometry, raw abundances were processed as described in figure legends.

### Isolation of CENP-T-based kinetochore-like particles from HeLa cells

HeLa cells with doxycycline inducible expression of GFP-I3-01 or GFP-CENP-T^1–242^–I3-01 were cultured in 15-cm tissue culture plates. After cells reached 70-90% confluence, expression of oligomers was induced by addition of 1 μg/ml doxycycline (Sigma-Aldrich, D9891) for 24 hrs. Mitotic cells were harvested by shaking off and gently rinsing with a pipette. Harvested cells were pelleted by centrifugation at 1,000 *g* and washed in DPBS buffer (Corning, 21-031-CV). Cells were resuspended in Lysis Buffer (50 mM HEPES pH 7.2, 2 mM MgCl_2_, 150 mM K-glutamate, 0.1 mM EDTA, 2 mM EGTA, 10% glycerol) and pelleted by centrifugation. Cell pellets containing ≈10^7^ cells were snap-frozen and stored in liquid nitrogen. Pellets of cells expressing GFP-I3-01 oligomers were prepared analogously except the mitotic cells were induced by adding 10 μM S-trityl-L-cysteine (STLC; Sigma-Aldrich, 164739) for 14 hrs.

GFP-CENP-T^1–242^–I3-01 and GFP-I3-01 oligomers were isolated from mitotic cell extracts prepared as in Tarasovetc, et al. 2021^25^. Briefly, one frozen cell pellet (∼100 µl) was resuspended in two volumes of ice-cold Lysis buffer supplemented with 0.1% IGEPAL (Sigma-Aldrich, I8896), 4 mM Mg-ATP, 2 mM DTT, protease inhibitors (0.2 mM 4-(2-Aminoethyl)-benzenesulfonylfluoride hydrochloride (AEBSF; Goldbiom, A-540-5), 10 μg/ml leupeptin (Roche, 11017128001), 10 μg/ml pepstatin (Roche, 11359053001), 10 μg/ml chymostatin (Sigma-Aldrich, C7268), Complete Mini EDTA free cocktail (Roche, 11836170001)), phosphatase inhibitors (100 ng/ml microcystin-LR (Enzo Life Sciences, ALX-350-012), 1 mM sodium pyrophosphate (Sigma-Aldrich, P8010), 2 mM sodium-beta-glycerophosphate (Santa Cruz Biotechnology, sc-220452), 100 nM sodium orthovanadate (Alfa Aesar, 81104-14), 5 mM sodium fluoride (Sigma-Aldrich, S6776), 120 nM okadaic acid (EMD Millipore, 495604), PhosSTOP cocktail (Roche, 04906845001)), and ATP regeneration system (10 mM phosphocreatine (Sigma-Aldrich, P7936), 0.45 mg/ml phospho-creatine kinase (Sigma-Aldrich, C3755)). Cells were ruptured by sonication using a Branson SFX150 Sonifier with a 3/32" microtip at 68% power for four cycles consisting of 15 s ON and 30 s OFF. During the entire procedure, the microcentrifuge tubes with cell suspension were kept in ice-cold water. Ruptured cells were treated with 1 U/μl OmniCleave endonuclease (Lucigen, OC7850K) for 5 min at 37°C to release the DNA-bound protein pool, and cells were sonicated for one more cycle. The suspension was centrifuged at 4,000 g for 15 min at 4°C, supernatant was collected and the oligomers were pelleted by ultracentrifugation at 280,000 g for 15 min at 4°C. Pellets were washed three times by gently adding and removing of 100 µl Lysis Buffer supplemented with all components described above, CENP-T-based kinetochore-like particles were resuspended in 50 µl of the same buffer, immediately aliquoted and snap-frozen in liquid nitrogen for storage at -80°C.

### Determining the size of GFP-containing oligomers *in vitro*

Experiments were performed using a Nikon Eclipse Ti microscope equipped with 1.49xNA TIRF 100xOil objective. A CUBE 488-nm 100 mW diode laser (Coherent) provided excitation to visualize GFP-tagged proteins in total internal reflection fluorescence (TIRF) mode. A CUBE 640-nm 50 mW diode laser and a CUBE 561-nm 100 mW diode laser (Coherent) provided excitation for microtubules polymerized from tubulins labeled with HiLyte647 or rhodamine. Images were acquired with an Andor iXon3 EMCCD camera and analyzed using Fiji software^68^. The size of oligomers with the GFP-tagged proteins was determined by measuring their fluorescence intensity and dividing by the intensity of one GFP molecule, which was determined under identical imaging conditions.

First, to determine brightness of single GFP molecule, a flow chamber was incubated for 1 min with 100 pM recombinant 6His-GFP, which was purified using a protocol for His-tagged proteins in Rago, et al. 2015 in Mg-BRB80 buffer ^14^ (80 mM K-1,4-Piperazinediethanesulfonic acid (K-PIPES) pH 6.9, 4 mM MgCl₂, 1 mM EGTA), washed and sealed with VALAP (1:1:1 vaseline/lanolin/paraffin). Bleaching of individual GFP spots (Supplementary Figure 2E) was captured for 1 min under TIRF illumination with the 20% laser power and the following settings for Andor iXon3 camera: 1 MHz readout speed, gain 5.0x, EM gain 50, 300 ms exposure time. To take into account an unevenness of laser illumination, images of GFP molecules were normalized on the laser intensity profile, which was generated by averaging >100 images of randomly selected fields with GFP molecules at high density (1 nM GFP). An integral intensity of individual GFP-molecules as a function of illumination time was measured in a circle area with the radius 3 pixels, generating individual photo-bleaching curves. Background intensity was measured in the same size area located near each GFP spot; individual background values were averaged for all examined spots and the resultant curve was subtracted from the individual photo-bleaching curves. Further processing, such as smoothing with the sliding window of 4 points and curves alignment, was carried out, as in Volkov, et al. 2014^71^. Individual photo-bleaching curves were combined to build a histogram, in which the non-zero peak was fitted with Gaussian function to represent the mean value of single molecule intensity (Supplementary Figure 2E).

Second, GFP-labeled oligomers isolated from mitotic cells were diluted 1,000-40,000 times in Mg-BRB80 buffer, flowed into the chamber and incubated for 5 min to immobilize them on the coverslip. Images were captured using the same camera settings as for single GFP molecules except the EM gain was decreased to 10. A linearity of EM gain settings was confirmed in separate experiments. Images of GFP-labeled oligomers were normalized on the laser intensity profile, and the oligomers were automatically selected using Fiji “Find Maxima” plugin with 5,800 prominence level, which excluded small GFP-labelled oligomers. Finally, the integral intensity of individual oligomers and corresponding background were measured in a circle area with the radius 6 pixels. After background subtraction, number of GFP-tagged molecules per oligomer was calculated as a ratio of intensity of this oligomer divided by average intensity of single GFP molecule and multiplied by 5 to take into account difference in EM Gain settings.

### Assays with stabilized microtubules *in vitro*

Tubulin for microtubules was purified from cow brains by thermal cycling and chromatography^72^, and labeled with HiLyte647 (HiLyte Fluor 647 succinimidyl ester; Anaspec, 81256), rhodamine (5-(and-6)-carboxytetramethylrhodamine succinimidyl ester; Invitrogen, C1171) or biotin (D-biotin succinimidyl ester; Invitrogen, B1513), as in Hyman, et al. 1991^73^. Taxol-stabilized fluorescent microtubules were prepared, as in Chakraborty, et al. 2018 from a mixture of unlabeled and HiLyte647-labeled tubulin^74^ (9:1, total tubulin concentration 100 μM). Custom-made flow chambers were assembled with silanized coverslips (22×22 mm) using spacers made from two strips of double-sided sticky tape, as in Chakraborty, et al. 2018^74^. Solutions were perfused with syringe pump (New Era Pump Systems, cat # NE-4000) and all experiments were carried out at 32°C. To immobilize taxol-stabilized microtubules, anti-tubulin antibodies (Serotec, MCA2047) were flowed into the chamber and the coverslip was blocked with 1% Pluronic F-127 (Sigma-Aldrich, CP2443) prior to introducing fluorescently labeled microtubules in Mg-BRB80 buffer supplemented with 7.5 μM taxol. Oligomers assembled on GFP-labeled clusters were then added in Imaging Buffer (Mg-BRB80 supplemented with 10 mM DTT, 7.5 μM taxol, 5 mM Mg-ATP, 4 mg ml^-1^ bovine serum albumin (BSA; Sigma-Aldrich, A7638), 0.1 mg ml^-1^ casein (Sigma-Aldrich, C5890), 0.1 mg ml^−1^ glucose oxidase (Sigma-Aldrich, G2133), 20 μg ml^−1^ catalase (Sigma-Aldrich, C40) and 6 mg ml^−1^ glucose (Sigma-Aldrich, G8270), incubated for 5 min, and then GFP and microtubule images were collected in TIRF mode. To allow quantitative comparison of the level of microtubule decoration by different oligomers, a care was taken to prepare solutions of clusters at similar concentration. To estimate concentration of isolated oligomers, thawed cluster suspensions were diluted 1,000-40,000-fold in Mg-BRB80 buffer and allowed to bind to the plasma cleaned coverslips for 5 min. Images of at least 10 different microscopy fields were collected, and the number of clusters per field was determined. Concertation of oligomers was calculated as the average number of GFP-labeled oligomers per imaging field multiplied by the dilution factor. Concertation of T-particles was 10-20 times lower than preparations with control GFP-oligomers, so the latter were diluted additionally to compensate for this difference. To quantify microtubule decoration with different oligomers, 10-20 rhodamine-microtubules per imaging field were selected in the rhodamine channel. Then, the number of GFP-labeled oligomers colocalizing with these microtubules was determined in the GFP channel by automatic selection with Fiji “Find Maxima” plugin with 5,800 prominence level.

### Assays with dynamic microtubules *in vitro*

Microtubule seeds were prepared, as in ^74^ from a mixture of unlabeled, rhodamine- and biotin-labeled tubulins (8:1:1, total tubulin concentration 5 μM) supplemented with 1 mM GMPCPP (Jena Bioscience, NU-405L). A flow chamber was prepared as for assays with taxol-stabilized microtubules, but the coverslip was coated with 5 μM neutravidin to assist immobilization of the biotin-containing microtubule seeds. Imaging Buffer supplemented with 1 mM Mg-GTP, a mixture of unlabeled and HiLyte647-labeled tubulin (8:2, total tubulin concentration 5 μM) and up to 0.3% methyl cellulose was flowed using the pump. Microtubule growth was observed for 5 min, and then CENP-T-based kinetochore-like particles were flowed into the chamber. Imaging was carried out in TIRF mode switching between 488-nm and 640-nm lasers with 300 ms exposure using stream acquisition at 12 frames per min. To analyze tracking, a microtubule visible via Hilyte647 fluorescence was fitted with a straight line (5 pixels width) using Fiji software, and the kymograph in microtubule and GFP-channels was prepared along this line. Kymographs with a bright GFP-dot at the end of microtubule relative to lattice were scored as the tip-tracking events. Polymerization and depolymerization microtubule rates were determined from the slopes of the corresponding kymographs.

### Immunostaining of isolated CENP-T-based kinetochore-like particles

Immunostaining was performed on CENP-T-based kinetochore-like particles bound to taxol-stabilized microtubules or oligomers immobilized on the coverslips functionalized by 10-min incubation with 20 µg ml^-1^ anti-S-tag antibodies (Abcam, ab87840) and blocked with 1% Pluronic F-127. Oligomers isolated from HeLa cells were allowed to adsorb onto the coverslip for 20 min. Chambers incubated with 3.5% paraformaldehyde or with no added fixative produced similar results, so these data were combined. Chambers were washed with Blocking Buffer (BRB80 buffer supplemented with 2 mM DTT, 4 mg ml^-1^ BSA and 0.5 mg ml^-1^ casein). Anti-Ndc80 antibodies ^49^ diluted at 25 μg ml^-1^ in Blocking Buffer were incubated for 15 min, followed by Alexa647–conjugated anti-rabbit antibodies (1:100, Thermo Fisher Scientific, A21245) for 15 min, and washed with Imaging Buffer. To analyze level of Ndc80 recruitment, GFP-labeled oligomers were selected in GFP-channel and the corresponding level of associated Ndc80 was measured as integral intensity in Alexa647 channel.

### Single molecule assay to measure CENP-T-NDC80 binding *in vitro*

Flow chambers were prepared as described above in the section “Assays with stabilized microtubules in vitro”. The surface of the coverslip was activated by incubation with 20 µg/ml anti-S-tag antibodies (Abcam) diluted in BRB80 buffer for 10 min. The coverslip was blocked with 1% Pluronic F-127. Then, GFP-CENP-T1-242/3D or GFP oligomers were introduced to the flow chamber. The specimen on the microscope stage was maintained at 32°C. The chamber was incubated for 20 min to allow immobilization of oligomers onto the coverslip. After immobilization oligomers were transferred to Imaging Buffer. Five images of the same field with GFP-tagged oligomers were collected for subsequent quantifications of their initial fluorescent intensity, which corresponds to the quantity of GFP-CENP-T1-242/3D or GFP molecules per oligomer. The oligomers were then bleached with a laser at 100% power for 30 s. Five images of the same field with oligomers were collected after bleaching to evaluate the efficiency of bleaching and the remaining GFP intensity of oligomers. Next, 100 μl of 100 nM GFP-tagged NDC80 in Imaging Buffer was introduced to the chamber using syringe pump at speed 900 μl/min. After 10 minutes, NDC80 was washed out from the chamber using 300 μl of Imaging Buffer perfused at speed 900 μl/min. Chamber was incubated for additional 10 min, and five images of the oligomers were collected to record recruitment of GFP-tagged NDC80.

The images were analyzed using Fiji^68^. First, the image sequence was corrected on the stage drift using “Manual drift correction” plugin. Then, the GFP intensity was measured in area surrounding the oligomer (8 pixels radius). Brightness of the same size area located near each oligomer was subtracted to minimize variability in background intensity. The intensities of individual oligomers during different stages of experiment were averaged between five frames. Final fluorescence intensity from GFP-tagged NDC80 was normalized on initial intensity from GFP-CENP-T1-242/3D or GFP oligomers. Resulting values represent average number of GFP-tagged NDC80 molecules per GFP-CENP-T1-242/3D or GFP molecule in oligomer.

The experiment with monomeric GFP-CENP-T^1–242/3D^ was done analogously with several modifications. To obtain a field with evenly dispersed molecules, 0.25 nM GFP-CENP-T^1–242/3D^ was used. GFP-CENP-T^1–242/3D^ molecules were not photobleached, to avoid confusion between detached GFP-CENP-T^1–242/3D^ molecules or those did not bind GFP-tagged NDC80. One frame was collected initially, and one was collected after NDC80 binding to avoid photobleaching. The probability of photobleaching was estimated from a photobleaching curve to be 4% over the 0.6 second exposure (Supplementary Figure 7G). Smaller areas (3 pixels radius) surrounding GFP-CENP-T^1–242/3D^ dots were used to measure their fluorescent intensity. To confirm, that GFP-CENP-T^1–242/3D^ is monomeric, distribution of their initial fluorescent intensities was normalized to the fluorescence of one GFP-molecule (Supplementary Figure 7C).

### Quantification and statistical analyses

Fiji/ImageJ (NIH) was used for image manipulation^68^. Statistical tests were performed in Graphpad Prism as described in figure legends.

## Acknowledgements

We thank Jimmy Ly and Steve Bell for feedback on the manuscript, we thank Océane Marescal for feedback on the manuscript and technical assistance, and we thank the members of the Cheeseman and Grishchuk labs for feedback throughout the process. This work was supported by grants to IMC from the NIH/NIGMS (R35GM126930), including diversity supplement funding for GBS, and the Gordon and Betty Moore Foundation, a grant to ELG from NIH/NIGMS (R35-GM141747), and grants to both IMC and ELG from the American Cancer Society Theory Lab Collaborative Grant (TLC-20-117-01-TLC) and NSF (2029868).

## Data Availability

The data that support the findings of this study are available from the corresponding author upon reasonable request.

---

**Supplementary Figure 1:**
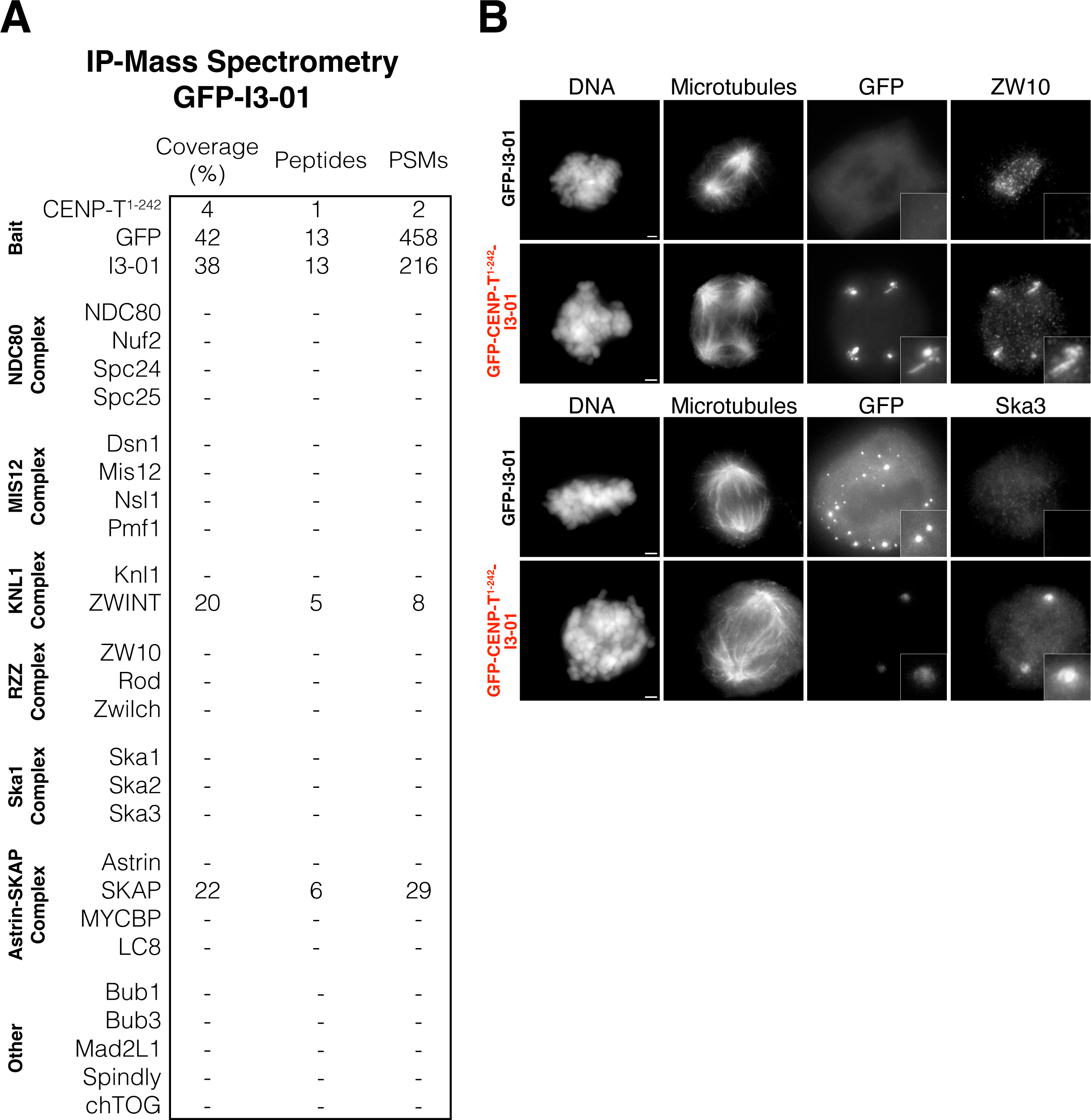
CENP-T^1–242^ oligomers recruit additional outer kinetochore proteins, but control oligomers do not. (A) Kinetochore and kinetochore-associated proteins detected in immuno-precipitation mass spectrometry of GFP-I3-01 control oligomers. (B) Co-localization of outer kinetochore proteins with GFP-CENP-T^1–242^-I3-01 oligomers by immunofluorescence. Image sets using the same antibody are scaled the same.

**Supplementary Figure 2.**
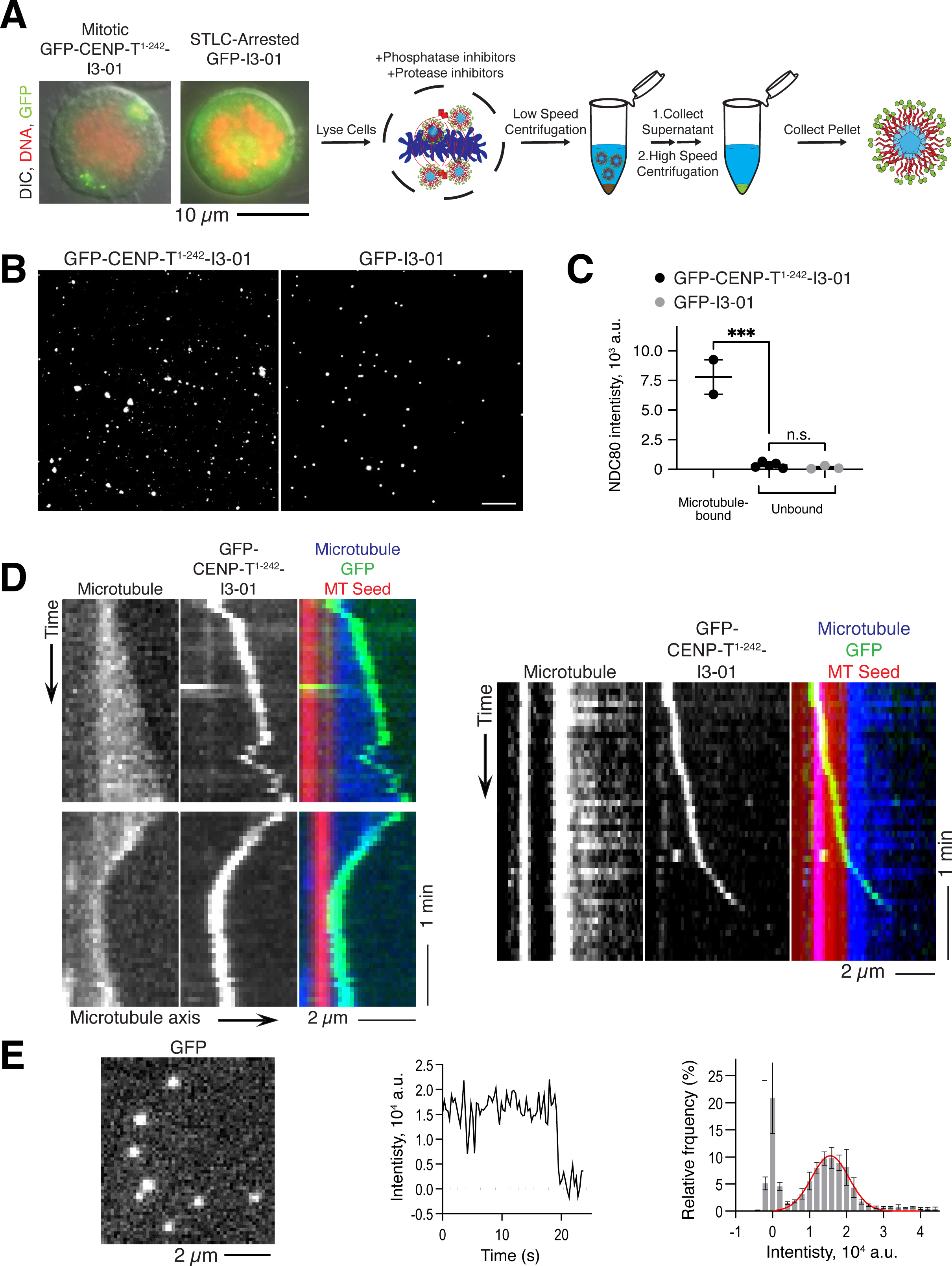
Characterization of GFP CENP-T^1–242^ and control GFP oligomers isolated from HeLa cells. (A) Workflow showing the key steps to isolate GFP-CENP-T^1–242^-I3-01 and GFP-I3-01 oligomers from mitotic cells. Left image shows representative HeLa cells expressing GFP-CENP-T^1–242^or GFP oligomers. Images are merges of DIC (grey scale), DAPI-stained DNA (red), and GFP (green). Expression of GFP-CENP-T^1–242^-I3-01 caused cells to arrest in mitosis. Cells expressing GFP-I3-01 were arrested in mitosis with the Eg5 inhibitor S-Trityl-L-Cysteine (STLC; see Methods for details). (B) Representative fluorescence microscopy images of the indicated GFP-labelled oligomers immobilized on coverslips; the same microscopy settings and linear image adjustments were used to allow direct comparison of oligomer brightness. (C) The graph shows NDC80 levels associated with CENP-T^1–242^-oligomers and GFP control oligomers bound to taxol-stabilized microtubules or settled on the coverslip (not bound to a microtubule). NDC80 was visualized using antibodies against the NDC80 complex. Note that only CENP-T-containing oligomers localize to microtubules, and that these complexes exhibit robust NDC80 association. Both types of oligomers that had settled on the coverslips contained little NDC80, suggesting that it dissociates from some CENP-T^1–242^ oligomers during the isolation procedure. Each point represents the median value from independent experiment with n>100 oligomers, lines and error bars show mean ± SEM from 2-5 independent experiments. P-values were calculated with unpaired two-tailed t-tests: n.s., p>0.05; ***, p<0.001. For more detailed statistics, see Source data. (D) Additional kymographs illustrating complex motions of CENP-T^1–242^ oligomers on dynamic microtubules, see legend to Figure 3F for more details. Top left: CENP-T^1–242^ oligomer moves along the microtubule wall to reach the plus-end, tracks the polymerizing plus-end, then diffuses along the microtubule wall. Bottom left: CENP-T^1–242^ oligomer remains associated with the microtubule end as it depolymerizes, then continues to track the end when it reverts to polymerization. Right: CENP-T^1–242^ oligomer moves unidirectionally on microtubule wall toward the growing plus-end. Note the different velocities on the GMPCPP-containing seed (red) and the GDP-containing lattice (blue; 0.7 μm/min vs. 3 μm/min). Faster motion on GDP vs. GMPCPP microtubule lattice has been reported for purified CENP-E kinesin ^38^, indirectly suggesting that some of the isolated CENP-T^1–242^ oligomers may also harbor CENP-E kinesin. (E) Quantification of the number of GFP molecules. Left: representative image of a microscope field with single GFP molecules immobilized on plasma-cleaned coverslip. Middle: Example photobleaching curve for a single molecule of GFP. Right: Histogram of integral intensities collected from 60 bleaching GFP dots from 3 independent experiments. Red line is fit to Gaussian function. Peak value of 1.56 ± 0.04 x 10^4^ a.u. corresponds to the integral intensity of a single GFP fluorophore under our imaging conditions. This intensity was used to estimate number of GFP fluorophores in oligomers and complexes, see Methods for details.

**Supplementary Figure 3:**
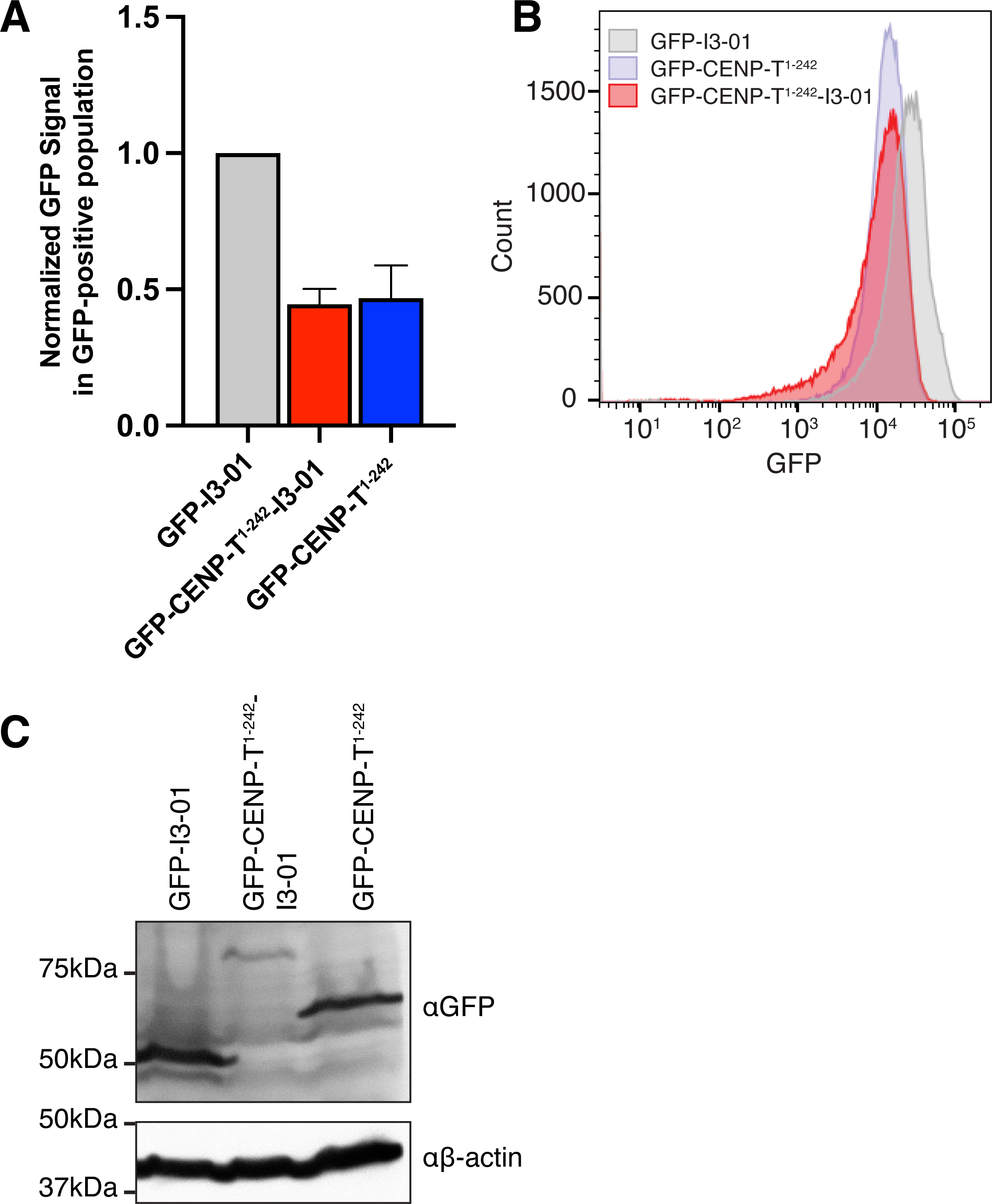
CENP-T^1–242^ oligomers, CENP-T^1–242^ monomers, and control oligomers are expressed at comparable levels. (A) Normalized GFP signals from GFP-positive cells analyzed for DNA content in Figure 4E as measured by flow cytometry. Averaged from three experiments (mean ± SD). The same cell lines were used for other assays with these three constructs. (B) Histograms showing the distribution of GFP expression levels in cells from cell line in (A) as measured by flow cytometry. (C) Anti-GFP and Anti-β-Actin western blots of cells expressing GFP-CENP-T^1–242^-I3-01, GFP-CENP-T^1–242^, and GFP-I3-01. β-Actin was used as a loading control.

**Supplementary Figure 4:**
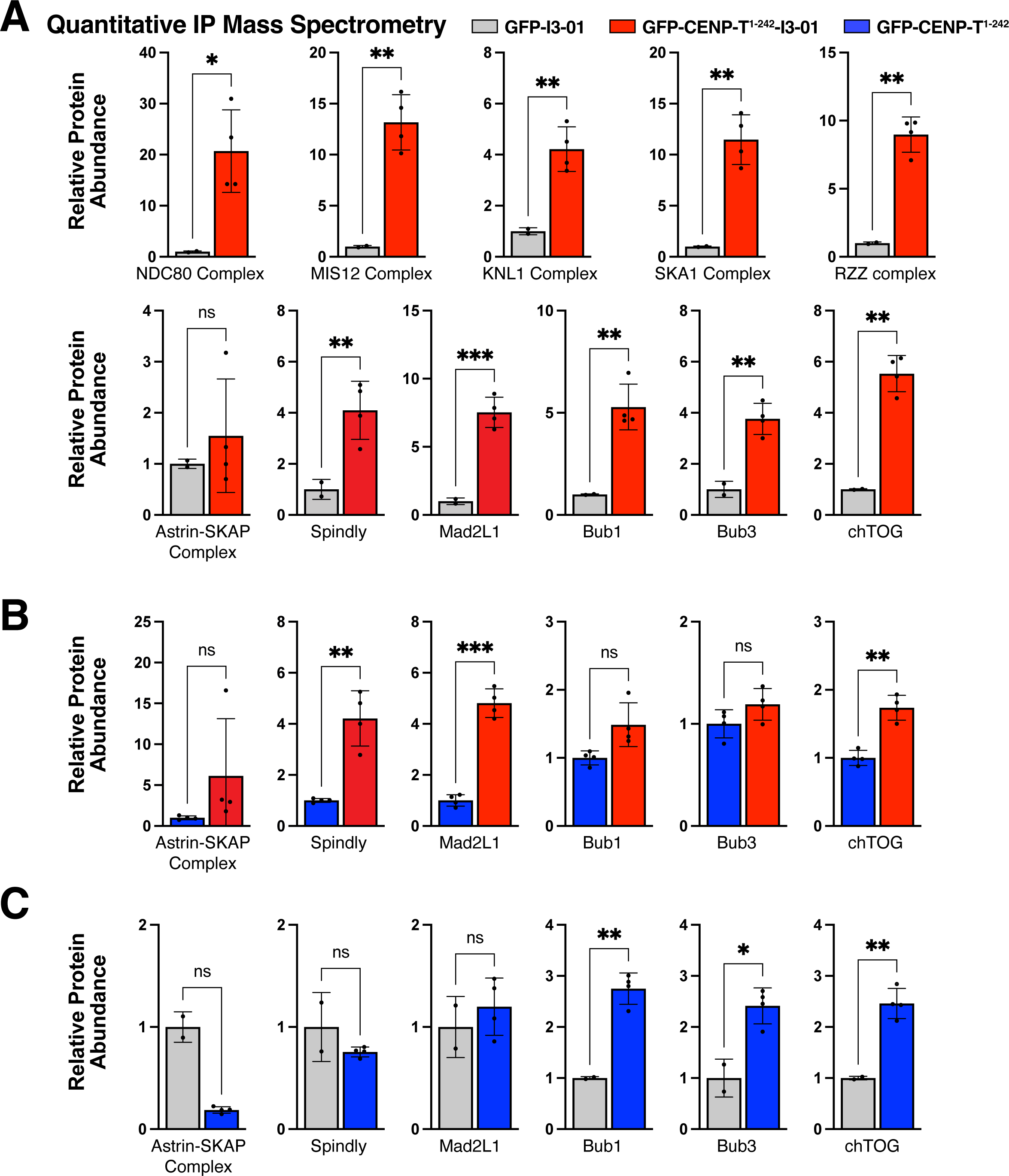
CENP-T^1–242^ oligomers recruit kinetochore-associated proteins and spindle assembly checkpoint proteins more robustly than monomeric CENP-T^1–242^. (A) Comparison of protein co-immuno-precipitation by CENP-T^1–242^ oligomers and control GFP oligomers by TMT-based quantitative immune-precipitation mass spectrometry. Each point represents a biological replicate from a single multiplexed mass spectrometry run. P-values were calculated using Welch’s two-tailed t-test. ns = p > 0.05; * = p < 0.05; ** = p <0.01; *** = p <0.001. Raw abundances of proteins in each sample were normalized to the abundances of GFP and I3-01 peptides, then the normalized abundances for the designated proteins or complexes were expressed as multiples of the normalized abundance in the GFP-I3-01 sample. Complex abundances were obtained by calculating the sum of abundances of the complex components. (B) Comparison of protein co-immuno-precipitation by CENP-T^1–242^ oligomers and CENP-T^1–242^ monomers by TMT-based quantitative immune-precipitation mass spectrometry. Each point represents a biological replicate from a single multiplexed mass spectrometry run. Raw abundances of proteins in each sample were normalized and added as described in Figure 4G. P-values were calculated as described in (A). (C) Comparison of protein co-immuno-precipitation by control GFP oligomers and CENP-T^1–242^ monomers by TMT-based quantitative immuno-precipitation mass spectrometry. Each point represents a biological replicate from a single multiplexed mass spectrometry run. Raw abundances of proteins in each sample were normalized and added as described in Figure 4F. P-values were calculated as described in (A).

**Supplementary Figure 5:**
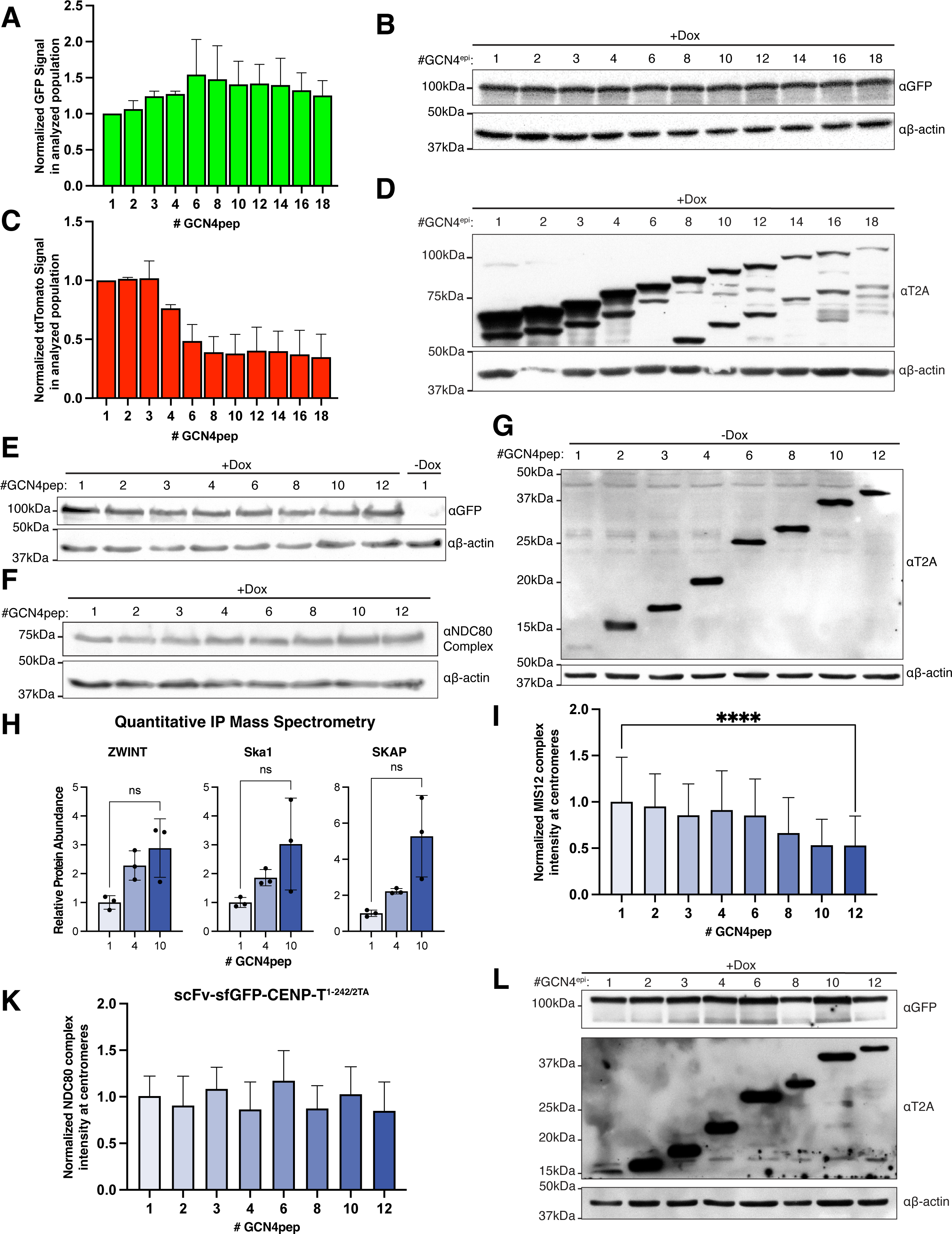
Construct expression from SunTag cell lines, additional SunTag mass spectrometry and centromere depletion data. (A) Normalized GFP signals from cells analyzed for DNA content in Figure 5C as measured by flow cytometry. Averaged from three experiments (mean ± SD). (B) Anti-GFP and Anti-β-Actin western blots of cell lines expressing scFv-sfGFP-CENP-T^1–242^ alongside tdTomato-tagged scaffolds with different numbers of GCN4pep. Anti-T2A antibody binds to the C-terminus of the scaffolds. β-Actin was used as a loading control. These cell lines were used in Figure 5C. (C) Normalized tdTomato signals from cells analyzed for DNA content in Figure 5C as measured by flow cytometry. Averaged from three experiments (mean ± SD) (D) Anti-T2A and Anti-β-Actin western blots of cell lines expressing scFv-sfGFP-CENP-T^1–242^ alongside tdTomato-tagged scaffolds with different numbers of GCN4pep. β-Actin was used as a loading control. These cell lines were used in Figure 5C. (E) Anti-GFP and Anti-β-Actin western blots of cell lines expressing scFv-sfGFP-CENP-T^1–242^ alongside myc-tagged scaffolds with different numbers of GCN4pep. 1xGCN4pep cell line without dox-treatment was used as a negative control, and β-Actin was used as a loading control. These cell lines were used in Figures 5E and 5F and in Supplementary Figure 5I. (F) Anti-NDC80 Complex and Anti-β-Actin western blots of cell lines expressing scFv-sfGFP-CENP-T^1–242^ alongside myc-tagged scaffolds with different numbers of GCN4pep. β-Actin was used as a loading control. These cell lines were used in Figures 5E and 5F and in Supplementary Figures 5I. (G) Anti-T2A and Anti-β-Actin western blots of cell lines expressing scaffolds with different numbers of GCN4pep. scFv-sfGFP-CENP-T^1–242^ expression was not induced in these experiments. Anti-T2A antibody binds to the C-terminus of the scaffolds. β-Actin was used as a loading control. These cell lines were used in Figures 5E and 5F and in Supplementary Figure 5I. (H) Comparison of outer kinetochore protein co-immuno-precipitation by SunTag oligomers expressed with scaffolds with 1, 4, or 10 copies of GCN4pep by TMT-based quantitative immune-precipitation mass spectrometry. Each point represents a biological replicate from a single multiplexed mass spectrometry run. P-values were calculated using Welch’s two-tailed t-test. n.s. = p > 0.05. (I) Quantification of MIS12 levels at centromeres in cells expressing the scFv-sfGFP-CENP-T^1–242^ with scaffolds with different numbers of GCN4pep. Each bar represents the mean ± SD of MIS12 signal from cells expressing the designated construct. Values for each cell were calculated from the mean of the MIS12 signals of kinetochores in that cell. Before calculating the mean for a cell, the MIS12 signal of each kinetochore in the cell was normalized to anti-centromere antibody signal from that kinetochore. Overall means were calculated from pooled data from multiple experiments. 25-51 total cells were analyzed from 3 different experiments for each condition. Welch’s ANOVA test was performed to calculate P-value for the whole dataset: P < 0.0001. Welch’s t test was used to calculated P-value for 1 v. 12: p < 0.0001. (K) Quantification of NDC80 levels at centromeres in cells expressing scFv-sfGFP-CENP-T^1–242/2TA^ with scaffolds with different numbers of GCN4pep. Each bar represents the mean ± SD of NDC80 signal from cells expressing the designated construct. Analysis was performed as described in (I). 27-33 total cells were analyzed and pooled from 2 different experiments for each condition. Welch’s ANOVA test was performed to calculate P-value for the whole dataset: P < 0.0001. Welch’s t test was used to calculated P-value for 1 v. 12: p < 0.5. (L) Anti-GFP, Anti-T2A, and Anti-β-Actin western blots of cell lines expressing scaffolds with different numbers of GCN4pep alongside scFv-sfGFP-CENP-T^1–242^/2TA. Anti-T2A antibody binds to the C-terminus of the scaffolds. β-Actin was used as a loading control. These cell lines were used in in Supplementary Figure 5K.

**Supplementary Figure 6:**
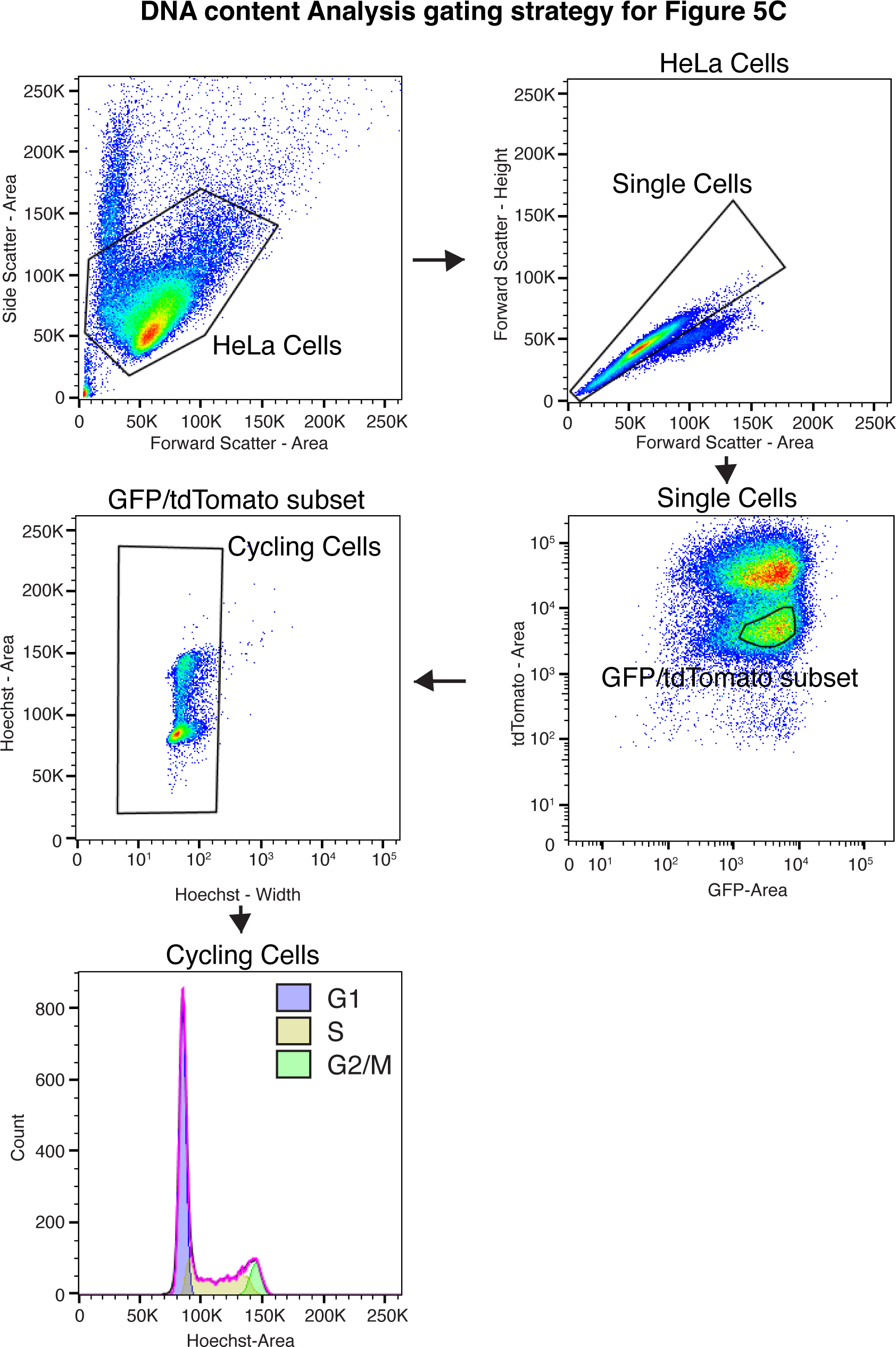
Flowcytometry gating strategy for DNA content analysis of CENP-T^1–242^ SunTag oligomers. Gating strategy to select the population of cells to be analyzed for DNA content analysis in Figure 5C. A similar gating strategy was used in Figure 4E without the tdTomato-A parameter.

**Supplementary Figure 7.**
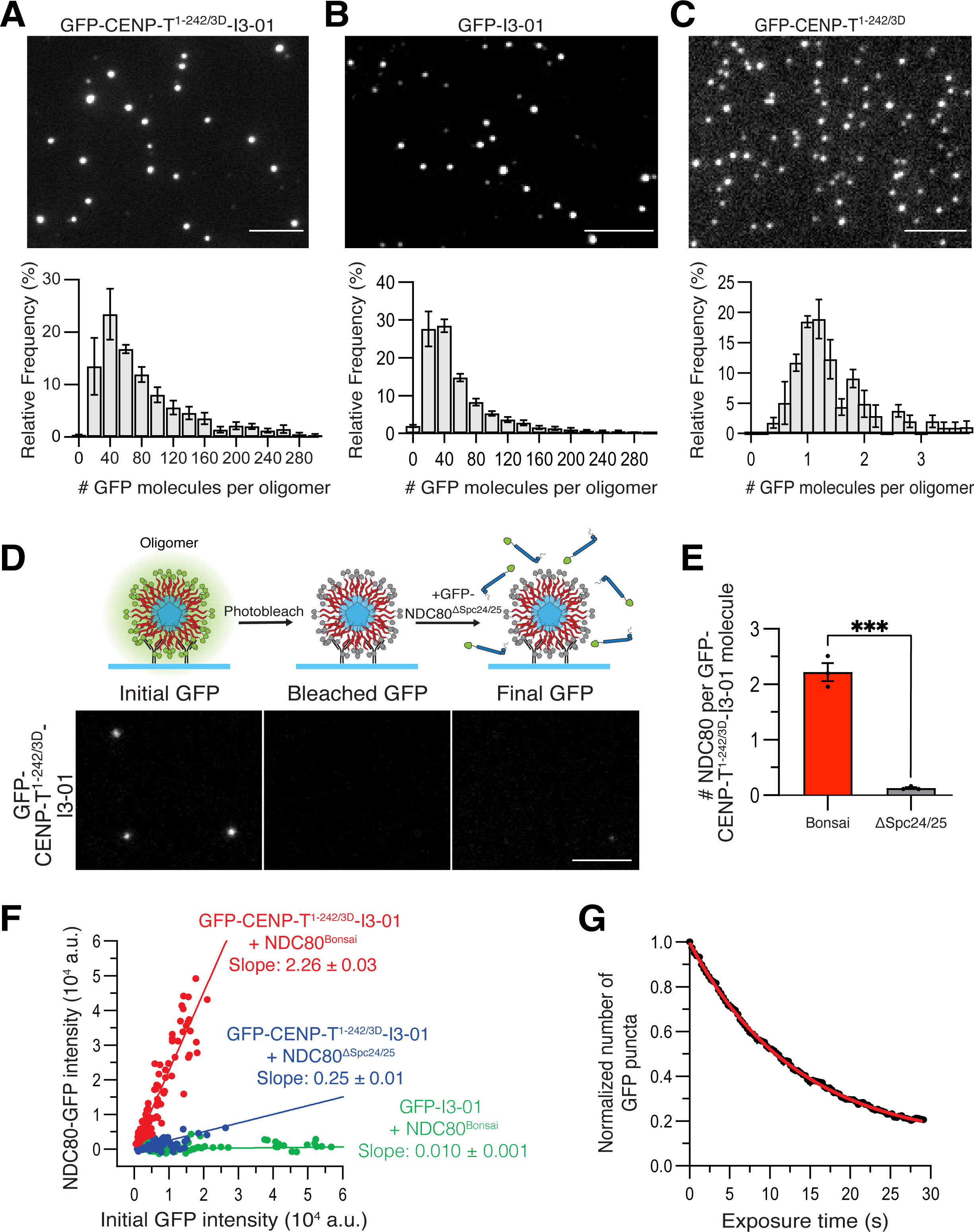
Additional fluorescence intensity quantifications for *in vitro* CENP-T-NDC80 binding assay using recombinant oligomers and NDC80 proteins. (A) Top: Representative images of purified recombinant GFP-CENP-T^1–242/3D^-I3-01 oligomers attached to a coverslip. Bottom: histogram showing the distribution of the number of GFP molecules per oligomer as a percentage of the total number of examined oligomers: 66 ± 10 GFP molecules (mean ± SEM of the distribution). Each bar represents the mean ± SEM from N=3 independent measurements, each with n>198 oligomers. (B) Same as panel A but for GFP-I3-01 control oligomers: 44 ± 4 GFP molecules (mean ± SEM of the distribution). Each bar represents the mean ± SEM from N=3 independent measurements, each with n>878 oligomers. (C) Same as panel A but for monomeric GFP-CENP-T^1–242/3D^ molecules: 1.23 ± 0.05 molecules (mean ± SEM of the distribution). Each bar represents the mean ± SEM from N=3 independent measurements, each with n>31 examined foci. Note that foci with several GFP molecules are rare and likely represent two or more molecules localizing close together. (D) Control experiment with NDC80^ΔSpc^^24/25^. Top: diagram of the experimental workflow. See Figure 6C legend for details. Bottom: representative images of GFP-CENP-T^1–242/3TD^-I3-01 oligomers before and after interaction with 100 nM of GFP-tagged NDC80^ΔSpc^^24/25^. (E) Efficiency of NDC80^Bonsai^ or NDC80^ΔSpc^^24/25^ recruitment to GFP-CENP-T^1–242/3D^-I3-01 oligomers. Bars are mean ± SEM from 3 independent experiments. Each point is the median result from independent trials with at least 12 oligomers analyzed. Data for GFP-CENP-T^1–242/3D^-I3-01 oligomer is duplicated from Figure 6D. P-values were calculated by unpaired two-tailed t-test: *** = p<0.001. (F) Graphical illustration of the stoichiometry of binding. Plot shows final GFP signal intensity as function of initial GFP signal intensity for individual oligomers in experiments using the indicated oligomers and NDC80 complexes. Each point represents the measurement for one oligomer from N=3 independent repeats per data set. Data are fitted to linear functions. The slopes (± standard fitting error) correspond to the number of NDC80 molecules recruited per GFP-containing monomer for each combination of oligomer and NDC80 complex. (G) Photobleaching curve taken with identical microscope settings to those used for experiments with GFP-CENP-T^1–242/3D^ (Figure 6B, E, F). The number of GFP puncta per imaging field at each time point was normalized to the number at t = 0. Data were fitted to an exponential decay function, which was used to estimate the probability of bleaching during imaging time. Each point represents the mean ± SEM from N=3 independent measurements.

## Supplementary Videos

**Supplementary Video 1. Kinetochore-like particle tracking a polymerizing microtubule end.** This video shows a dynamic microtubule (blue) growing from the coverslip-bound GMPCPP microtubule seed (red) in the presence of soluble tubulin and GFP-CENP-T^1–242^-I3-01 oligomers (green) isolated from mitotic HeLa cells. The isolated oligomer with associated proteins (i.e., kinetochore-like particle) binds to the end of a polymerizing microtubule and moves processively with the end as it polymerizes. This video plays at 7 fps and events are shown at 3.5 times actual speed.

**Supplementary Video 2. A kinetochore-like particle exhibits plus-end-directed processive movement.** A larger green particle remains stationary, while a smaller kinetochore-like particle moves processively and unidirectionally toward the growing microtubule plus-end. Initial motion on the GMPCPP-containing microtubule seed is slow, but the particle speeds up on the GDP-containing microtubule wall (see Supplementary Figure 2D for more details). These plus-end-directed motions were rarer than the other types of motion. This video is played at 7 fps and events are shown at 3.5 times actual speed.

**Supplementary Video 3. Kinetochore-like particle diffusing along a microtubule wall and tracking a depolymerizing microtubule end.** Video shows a dynamic microtubule interacting with two kinetochore-like particles. A larger particle associates with the microtubule wall and remains stationary for the duration of the video. A smaller particle lands on the microtubule wall at the site indicated with an arrow. The particle diffuses along the microtubule wall, then tracks the depolymerizing end of the microtubule. This video is played at 7 fps and events are shown at 3.5 times actual speed.

## References

1. Cheeseman, I.M. (2014). The kinetochore. Cold Spring Harb Perspect Biol. 10.1101/cshperspect.a015826.

2. Musacchio, A., and Desai, A. (2017). A Molecular View of Kinetochore Assembly and Function. Biology (Basel) 6, 5. 10.3390/biology6010005.

3. Hara, M., Ariyoshi, M., Sano, T., Nozawa, R., Shinkai, S., Onami, S., Jansen, I., Hirota, T., and Fukagawa, T. (2022). The centromere/kinetochore is assembled through CENP-C oligomerization. bioRxiv, 2022.08.17.504347. 10.1101/2022.08.17.504347.

4. Suzuki, A., Badger, B.L., and Salmon, E.D. (2015). A quantitative description of Ndc80 complex linkage to human kinetochores. Nat Commun. 10.1038/ncomms9161.

5. Brinkley, B.R., and Cart Wright, J. (1971). Ultrastructural analysis of mitotic spindle elongation in mammalian cells in vitro. Direct microtubule counts. J Cell Biol 50, 416–431. 10.1083/JCB.50.2.416.

6. Zhou, K., Gebala, M., Woods, D., Sundararajan, K., Edwards, G., Krzizike, D., Wereszczynski, J., Straight, A.F., and Luger, K. (2022). CENP-N promotes the compaction of centromeric chromatin. Nature Structural & Molecular Biology 2022 29:4 29, 403–413. 10.1038/s41594-022-00758-y.

7. Xia, S., Chen, Z., Shen, C., and Fu, T.M. (2021). Higher-order assemblies in immune signaling: supramolecular complexes and phase separation. Protein Cell 12, 680. 10.1007/S13238-021-00839-6.

8. Wu, H. (2013). Higher-order assemblies in a new paradigm of signal transduction. Cell 153, 287–292. 10.1016/j.cell.2013.03.013.

9. Wu, H., and Fuxreiter, M. (2016). The Structure and Dynamics of Higher-Order Assemblies: Amyloids, Signalosomes, and Granules. Cell 165, 1055–1066. 10.1016/j.cell.2016.05.004.

10. Banani, S.F., Lee, H.O., Hyman, A.A., and Rosen, M.K. (2017). Biomolecular condensates: organizers of cellular biochemistry. Nature Reviews Molecular Cell Biology 2017 18:5 18, 285–298. 10.1038/nrm.2017.7.

11. Marzahn, M.R., Marada, S., Lee, J., Nourse, A., Kenrick, S., Zhao, H., Ben-Nissan, G., Kolaitis, R.-M., Peters, J.L., Pounds, S., et al. (2016). Higher-order oligomerization promotes localization of SPOP to liquid nuclear speckles. EMBO J 35, 1254–1275. 10.15252/EMBJ.201593169.

12. Navarro, A.P., and Cheeseman, I.M. (2021). Kinetochore assembly throughout the cell cycle. Semin Cell Dev Biol 117, 62–74. 10.1016/J.SEMCDB.2021.03.008.

13. Gascoigne, K.E., and Cheeseman, I.M. (2013). CDK-dependent phosphorylation and nuclear exclusion coordinately control kinetochore assembly state. Journal of Cell Biology 201, 23–32. 10.1083/jcb.201301006.

14. Rago, F., Gascoigne, K.E., and Cheeseman, I.M. (2015). Distinct organization and regulation of the outer kinetochore KMN network downstream of CENP-C and CENP-T. Curr Biol 25, 671–677. 10.1016/j.cub.2015.01.059.

15. Hara, M., Ariyoshi, M., Okumura, E. ichi, Hori, T., and Fukagawa, T. (2018). Multiple phosphorylations control recruitment of the KMN network onto kinetochores. Nat Cell Biol 20, 1378–1388. 10.1038/s41556-018-0230-0.

16. Nishino, T., Rago, F., Hori, T., Tomii, K., Cheeseman, I.M., and Fukagawa, T. (2013). CENP-T provides a structural platform for outer kinetochore assembly. EMBO Journal 32, 424– 436. 10.1038/emboj.2012.348.

17. Huis In’t Veld, P.J., Jeganathan, S., Petrovic, A., Singh, P., John, J., Krenn, V., Weissmann, F., Bange, T., and Musacchio, A. (2016). Molecular basis of outer kinetochore assembly on CENP-T. Elife. 10.7554/eLife.21007.

18. Palladino, J., Chavan, A., Sposato, A., Mason, T.D., and Mellone, B.G. (2020). Targeted De Novo Centromere Formation in Drosophila Reveals Plasticity and Maintenance Potential of CENP-A Chromatin. Dev Cell 52, 379–394.e7. 10.1016/J.DEVCEL.2020.01.005.

19. Gascoigne, K.E., Takeuchi, K., Suzuki, A., Hori, T., Fukagawa, T., and Cheeseman, I.M. (2011). Induced ectopic kinetochore assembly bypasses the requirement for CENP-A nucleosomes. Cell. 10.1016/j.cell.2011.03.031.

20. Gascoigne, K.E., and Cheeseman, I.M. (2013). Induced dicentric chromosome formation promotes genomic rearrangements and tumorigenesis. Chromosome Research 21, 407– 418. 10.1007/s10577-013-9368-6.

21. Mckinley, K.L., and Cheeseman, I.M. (2015). The molecular basis for centromere identity and function. Nature Publishing Group 17, 16–29. 10.1038/nrm.2015.5.

22. van Hooser, A.A., Ouspenski, I.I., Gregson, H.C., Starr, D.A., Yen, T.J., Goldberg, M.L., Yokomori, K., Earnshaw, W.C., Sullivan, K.F., and Brinkley, B.R. (2001). Specification of kinetochore-forming chromatin by the histone H3 variant CENP-A. J Cell Sci 114, 3529– 3542.

23. Walstein, K., Petrovic, A., Pan, D., Hagemeier, B., Vogt, D., Vetter, I.R., and Musacchio, A. (2021). Assembly principles and stoichiometry of a complete human kinetochore module. Sci Adv 7. 10.1126/SCIADV.ABG1037/SUPPL_FILE/ABG1037_SM.PDF.

24. Yatskevich, S., Muir, K.W., Bellini, D., Zhang, Z., Yang, J., Tischer, T., Predin, M., Dendooven, T., McLaughlin, S.H., and Barford, D. (2022). Structure of the human inner kinetochore bound to a centromeric CENP-A nucleosome. Science (1979) 376, 844–852. 10.1126/SCIENCE.ABN3810/SUPPL_FILE/SCIENCE.ABN3810_MOVIES_S1_AND_S2.ZIP.

25. Tarasovetc, E. V., Allu, P.K., Wimbish, R.T., DeLuca, J.G., Cheeseman, I.M., Black, B.E., and Grishchuk, E.L. (2021). Permitted and restricted steps of human kinetochore assembly in mitotic cell extracts. Mol Biol Cell 32, 1241–1255. 10.1091/MBC.E20-07-0461/ASSET/IMAGES/LARGE/MBC-32-1241-G007.JPEG.

26. Nishino, T., Takeuchi, K., Gascoigne, K.E., Suzuki, A., Hori, T., Oyama, T., Morikawa, K., Cheeseman, I.M., and Fukagawa, T. (2012). CENP-T-W-S-X forms a unique centromeric chromatin structure with a histone-like fold. Cell 148, 487–501. 10.1016/j.cell.2011.11.061.

27. Hsia, Y., Bale, J.B., Gonen, S., Shi, D., Sheffler, W., Fong, K.K., Nattermann, U., Xu, C., Huang, P.S., Ravichandran, R., et al. (2016). Design of a hyperstable 60-subunit protein icosahedron. Nature 535, 136–139. 10.1038/nature18010.

28. Hori, T., Amano, M., Suzuki, A., Backer, C.B., Welburn, J.P., Dong, Y., McEwen, B.F., Shang, W.H., Suzuki, E., Okawa, K., et al. (2008). CCAN Makes Multiple Contacts with Centromeric DNA to Provide Distinct Pathways to the Outer Kinetochore. Cell 135, 1039– 1052. 10.1016/j.cell.2008.10.019.

29. Musacchio, A. (2015). The Molecular Biology of Spindle Assembly Checkpoint Signaling Dynamics. Current Biology 25, R1002–R1018. 10.1016/J.CUB.2015.08.051.

30. Monda, J.K., and Cheeseman, I.M. (2018). The kinetochore–microtubule interface at a glance. J Cell Sci 131, jcs214577. 10.1242/jcs.214577.

31. Kops, G.J.P.L., and Gassmann, R. (2020). Crowning the Kinetochore: The Fibrous Corona in Chromosome Segregation. Trends Cell Biol 30, 653–667. 10.1016/J.TCB.2020.04.006.

32. Cheeseman, I.M., and Desai, A. (2008). Molecular architecture of the kinetochore– microtubule interface. Nat Rev Mol Cell Biol 9, 33–46. 10.1038/nrm2310.

33. Grishchuk, E.L. (2017). Biophysics of Microtubule End Coupling at the Kinetochore. Prog Mol Subcell Biol 56, 397–428. 10.1007/978-3-319-58592-5_17/FIGURES/3.

34. Joglekar, A.P., Bloom, K.S., and Salmon, E.D. (2010). Mechanisms of force generation by end-on kinetochore-microtubule attachments. Curr Opin Cell Biol 22, 57. 10.1016/J.CEB.2009.12.010.

35. McIntosh, J.R., Grishchuk, E.L., Morphew, M.K., Efremov, A.K., Zhudenkov, K., Volkov, V.A., Cheeseman, I.M., Desai, A., Mastronarde, D.N., and Ataullakhanov, F.I. (2008). Fibrils Connect Microtubule Tips with Kinetochores: A Mechanism to Couple Tubulin Dynamics to Chromosome Motion. Cell 135, 322–333. 10.1016/j.cell.2008.08.038.

36. Craske, B., and Welburn, J.P.I. (2020). Leaving no-one behind: how CENP-E facilitates chromosome alignment. Essays Biochem 64, 313–324. 10.1042/EBC20190073.

37. Maiato, H., Gomes, A.M., Sousa, F., and Barisic, M. (2017). Mechanisms of Chromosome Congression during Mitosis. Biology 2017, Vol. 6, Page 13 6, 13. 10.3390/BIOLOGY6010013.

38. Gudimchuk, N., Vitre, B., Kim, Y., Kiyatkin, A., Cleveland, D.W., Ataullakhanov, F.I., and Grishchuk, E.L. (2013). Kinetochore kinesin CENP-E is a processive bi-directional tracker of dynamic microtubule tips. Nat Cell Biol 15, 1079–1088. 10.1038/ncb2831.

39. Cheeseman, I.M., Chappie, J.S., Wilson-Kubalek, E.M., and Desai, A. (2006). The Conserved KMN Network Constitutes the Core Microtubule-Binding Site of the Kinetochore. Cell 127, 983–997. 10.1016/J.CELL.2006.09.039.

40. Chakraborty, M., Tarasovetc, E. V., Zaytsev, A. V., Godzi, M., Figueiredo, A.C., Ataullakhanov, F.I., and Grishchuk, E.L. (2019). Microtubule end conversion mediated by motors and diffusing proteins with no intrinsic microtubule end-binding activity. Nature Communications 2019 10:1 10, 1–14. 10.1038/s41467-019-09411-7.

41. Cai, S., O’Connell, C.B., Khodjakov, A., and Walczak, C.E. (2009). Chromosome congression in the absence of kinetochore fibres. Nature Cell Biology 2009 11:7 11, 832– 838. 10.1038/ncb1890.

42. Hunt, A.J., and McIntosh, J.R. (1998). The dynamic behavior of individual microtubules associated with chromosomes in vitro. undefined 9, 2857–2871. 10.1091/MBC.9.10.2857.

43. Volkov, V.A., Huis In ’T Veld, P.J., Dogterom, M., and Musacchio, A. (2018). Multivalency of NDC80 in the outer kinetochore is essential to track shortening microtubules and generate forces. Elife 7, e36764. 10.7554/eLife.36764.

44. Powers, A.F., Franck, A.D., Gestaut, D.R., Cooper, J., Gracyzk, B., Wei, R.R., Wordeman, L., Davis, T.N., and Asbury, C.L. (2009). The Ndc80 Kinetochore Complex Forms Load-Bearing Attachments to Dynamic Microtubule Tips via Biased Diffusion. Cell 136, 865–875. 10.1016/J.CELL.2008.12.045.

45. Tanenbaum, M.E., Gilbert, L.A., Qi, L.S., Weissman, J.S., and Vale, R.D. (2014). A Protein-Tagging System for Signal Amplification in Gene Expression and Fluorescence Imaging. Cell 159, 635–646. 10.1016/J.CELL.2014.09.039.

46. Ciferri, C., Pasqualato, S., Screpanti, E., Varetti, G., Santaguida, S., dos Reis, G., Maiolica, A., Polka, J., de Luca, J.G., de Wulf, P., et al. (2008). Implications for Kinetochore-Microtubule Attachment from the Structure of an Engineered Ndc80 Complex. Cell 133, 427–439. 10.1016/J.CELL.2008.03.020.

47. Itzhak, D.N., Tyanova, S., Cox, J., and Borner, G.H.H. (2016). Global, quantitative and dynamic mapping of protein subcellular localization. Elife 5. 10.7554/ELIFE.16950.

48. Wiśniewski, J.R., Hein, M.Y., Cox, J., and Mann, M. (2014). A “Proteomic Ruler” for Protein Copy Number and Concentration Estimation without Spike-in Standards. Molecular & Cellular Proteomics 13, 3497–3506. 10.1074/MCP.M113.037309.

49. Schmidt, J.C., Arthanari, H., Boeszoermenyi, A., Dashkevich, N.M., Wilson-Kubalek, E.M., Monnier, N., Markus, M., Oberer, M., Milligan, R.A., Bathe, M., et al. (2012). The Kinetochore-Bound Ska1 Complex Tracks Depolymerizing Microtubules and Binds to Curved Protofilaments. Dev Cell 23, 968–980. 10.1016/J.DEVCEL.2012.09.012.

50. Zaytsev, A. V., Sundin, L.J.R., DeLuca, K.F., Grishchuk, E.L., and DeLuca, J.G. (2014). Accurate phosphoregulation of kinetochore-microtubule affinity requires unconstrained molecular interactions. Journal of Cell Biology 206, 45–59. 10.1083/JCB.201312107/VIDEO-3.

51. McKinley, K.L., and Cheeseman, I.M. (2015). The molecular basis for centromere identity and function. Nature Reviews Molecular Cell Biology 2015 17:1 17, 16–29. 10.1038/nrm.2015.5.

52. Nechemia-Arbely, Y., Miga, K.H., Shoshani, O., Aslanian, A., McMahon, M.A., Lee, A.Y., Fachinetti, D., Yates, J.R., Ren, B., and Cleveland, D.W. (2019). DNA replication acts as an error correction mechanism to maintain centromere identity by restricting CENP-A to centromeres. Nat Cell Biol 21, 743. 10.1038/S41556-019-0331-4.

53. Athwal, R.K., Walkiewicz, M.P., Baek, S., Fu, S., Bui, M., Camps, J., Ried, T., Sung, M.H., and Dalal, Y. (2015). CENP-A nucleosomes localize to transcription factor hotspots and subtelomeric sites in human cancer cells. Epigenetics Chromatin 8, 1–23. 10.1186/1756-8935-8-2/TABLES/7.

54. Altemose, N., Maslan, A., Smith, O.K., Sundararajan, K., Brown, R.R., Mishra, R., Detweiler, A.M., Neff, N., Miga, K.H., Straight, A.F., et al. (2022). DiMeLo-seq: a long-read, single-molecule method for mapping protein–DNA interactions genome wide. Nature Methods 2022 19:6 19, 711–723. 10.1038/s41592-022-01475-6.

55. Bodor, D.L., Mata, J.F., Sergeev, M., David, A.F., Salimian, K.J., Panchenko, T., Cleveland, D.W., Black, B.E., Shah, J. v., and Jansen, L.E.T. (2014). The quantitative architecture of centromeric chromatin. Elife 2014. 10.7554/ELIFE.02137.

56. Bhat, P., Honson, D., and Guttman, M. (2021). Nuclear compartmentalization as a mechanism of quantitative control of gene expression. Nature Reviews Molecular Cell Biology 2021 22:10 22, 653–670. 10.1038/s41580-021-00387-1.

57. Li, P., Banjade, S., Cheng, H.C., Kim, S., Chen, B., Guo, L., Llaguno, M., Hollingsworth, J. v., King, D.S., Banani, S.F., et al. (2012). Phase transitions in the assembly of multivalent signalling proteins. Nature 2012 483:7389 483, 336–340. 10.1038/nature10879.

58. Schmidt, H.B., and Görlich, D. (2016). Transport Selectivity of Nuclear Pores, Phase Separation, and Membraneless Organelles. Trends Biochem Sci 41, 46–61. 10.1016/J.TIBS.2015.11.001.

59. Schwarz-Romond, T., Fiedler, M., Shibata, N., Butler, P.J.G., Kikuchi, A., Higuchi, Y., and Bienz, M. (2007). The DIX domain of Dishevelled confers Wnt signaling by dynamic polymerization. Nature Structural & Molecular Biology 2007 14:6 14, 484–492. 10.1038/nsmb1247.

60. Li, J., McQuade, T., Siemer, A.B., Napetschnig, J., Moriwaki, K., Hsiao, Y.S., Damko, E., Moquin, D., Walz, T., McDermott, A., et al. (2012). The RIP1/RIP3 necrosome forms a functional amyloid signaling complex required for programmed necrosis. Cell 150, 339– 350. 10.1016/j.cell.2012.06.019.

61. Lu, A., Li, Y., Yin, Q., Ruan, J., Yu, X., Egelman, E., and Wu, H. (2015). Plasticity in PYD assembly revealed by cryo-EM structure of the PYD filament of AIM2. Cell Discovery 2015 1:1 1, 1–14. 10.1038/celldisc.2015.13.

62. Lu, A., and Wu, H. (2015). Structural mechanisms of inflammasome assembly. FEBS J 282, 435–444. 10.1111/FEBS.13133.

63. Park, H., Kim, N.Y., Lee, S., Kim, N., Kim, J., and Heo, W. Do (2017). Optogenetic protein clustering through fluorescent protein tagging and extension of CRY2. Nature Communications 2017 8:1 8, 1–8. 10.1038/s41467-017-00060-2.

64. Qian, K., Huang, C.L., Chen, H., Blackbourn, L.W., Chen, Y., Cao, J., Yao, L., Sauvey, C., Du, Z., and Zhang, S.C. (2014). A Simple and Efficient System for Regulating Gene Expression in Human Pluripotent Stem Cells and Derivatives. Stem Cells 32, 1230–1238. 10.1002/STEM.1653.

65. Bugaj, L.J., Choksi, A.T., Mesuda, C.K., Kane, R.S., and Schaffer, D. V. (2013). Optogenetic protein clustering and signaling activation in mammalian cells. Nature Methods 2013 10:3 10, 249–252. 10.1038/nmeth.2360.

66. Cong, L., Ran, F.A., Cox, D., Lin, S., Barretto, R., Habib, N., Hsu, P.D., Wu, X., Jiang, W., Marraffini, L.A., et al. (2013). Multiplex genome engineering using CRISPR/Cas systems. Science (1979) 339, 819–823. 10.1126/SCIENCE.1231143/SUPPL_FILE/PAPV2.PDF.

67. Wang, T., Birsoy, K., Hughes, N.W., Krupczak, K.M., Post, Y., Wei, J.J., Lander, E.S., and Sabatini, D.M. (2015). Identification and characterization of essential genes in the human genome. Science (1979) 350, 1096–1101. 10.1126/SCIENCE.AAC7041/SUPPL_FILE/WANG-SM.PDF.

68. Schindelin, J., Arganda-Carreras, I., Frise, E., Kaynig, V., Longair, M., Pietzsch, T., Preibisch, S., Rueden, C., Saalfeld, S., Schmid, B., et al. (2012). Fiji: an open-source platform for biological-image analysis. Nature Methods 2012 9:7 9, 676–682. 10.1038/nmeth.2019.

69. Stirling, D.R., Swain-Bowden, M.J., Lucas, A.M., Carpenter, A.E., Cimini, B.A., and Goodman, A. (2021). CellProfiler 4: improvements in speed, utility and usability. BMC Bioinformatics 22, 1–11. 10.1186/S12859-021-04344-9/FIGURES/6.

70. McQuin, C., Goodman, A., Chernyshev, V., Kamentsky, L., Cimini, B.A., Karhohs, K.W., Doan, M., Ding, L., Rafelski, S.M., Thirstrup, D., et al. (2018). CellProfiler 3.0: Next-generation image processing for biology. PLoS Biol 16. 10.1371/JOURNAL.PBIO.2005970.

71. Volkov, V.A., Zaytsev, A. V, and Grishchuk, E.L. (2014). Preparation of Segmented Microtubules to Study Motions Driven by the Disassembling Microtubule Ends. J. Vis. Exp 85, 51150. 10.3791/51150.

72. Miller, H.P., and Wilson, L. (2010). Preparation of Microtubule Protein and Purified Tubulin from Bovine Brain by Cycles of Assembly and Disassembly and Phosphocellulose Chromatography. Methods Cell Biol 95, 2–15. 10.1016/S0091-679X(10)95001-2.

73. Hyman, A.A., and Mitchison, T.J. (1991). Two different microtubule-based motor activities with opposite polarities in kinetochores. Nature 1991 351:6323 351, 206–211. 10.1038/351206a0.

74. Chakraborty, M., Tarasovetc, E. V, and Grishchuk, E.L. (2018). In vitro reconstitution of lateral to end-on conversion of kinetochore-microtubule attachments. Mitosis and Meiosis Part A, 1–21. 10.1016/bs.mcb.2018.03.018.

75. McKinley, K.L., and Cheeseman, I.M. (2017). Large-Scale Analysis of CRISPR/Cas9 Cell-Cycle Knockouts Reveals the Diversity of p53-Dependent Responses to Cell-Cycle Defects. Dev Cell. 10.1016/j.devcel.2017.01.012.

76. Cheeseman, I.M., Hori, T., Fukagawa, T., and Desai, A. (2008). KNL1 and the CENP-H/I/K complex coordinately direct kinetochore assembly in vertebrates. Mol Biol Cell 19, 587– 594. 10.1091/mbc.E07-10-1051.

77. Welburn, J.P.I., Grishchuk, E.L., Backer, C.B., Wilson-Kubalek, E.M., Yates, J.R., and Cheeseman, I.M. (2009). The Human Kinetochore Ska1 Complex Facilitates Microtubule Depolymerization-Coupled Motility. Dev Cell 16, 374–385. 10.1016/J.DEVCEL.2009.01.011.

